# NiCLIP: Neuroimaging contrastive language-image pretraining model for predicting text from brain activation images

**DOI:** 10.1101/2025.06.14.659706

**Authors:** Julio A. Peraza, James D. Kent, Thomas E. Nichols, Jean-Baptiste Poline, Alejandro de la Vega, Angela R. Laird

## Abstract

Predicting cognitive processes from brain activation maps has remained an open question within the neuroscience community for many years. Meta-analytic functional decoding methods aim to tackle this issue by providing a quantitative estimation of behavioral profiles associated with specific brain regions. Existing methods face intrinsic challenges in neuroimaging meta-analysis, particularly in consolidating textual information from publications, as they rely on limited metrics that do not capture the semantic context of the text. The combination of large language models (LLMs) with advanced deep contrastive learning models (e.g., CLIP) for aligning text with images has revolutionized neuroimaging meta-analysis, potentially offering solutions to functional decoding challenges. In this work, we present NiCLIP, a contrastive language-image pretrained model that predicts cognitive tasks, concepts, and domains from brain activation patterns. We leveraged over 23,000 neuroscientific articles to train a CLIP model for text-to-brain association. Evaluation of NiCLIP predictions revealed that performance is optimized when using full-text articles instead of abstracts, as well as a curated cognitive ontology with precise task-concept-domain mappings. Furthermore, domain-specific fine-tuned LLMs (e.g., BrainGPT models) show numerically similar performance to their base LLM counterparts. Our results indicated that NiCLIP accurately predicts cognitive tasks from group-level activation maps provided by the Human Connectome Project across multiple domains (e.g., emotion, language, motor) and precisely characterizes the functional roles of specific brain regions, including the amygdala, hippocampus, and temporoparietal junction. However, NiCLIP showed limitations with noisy subject-level activation maps. NiCLIP represents a significant advancement in quantitative functional decoding for neuroimaging, offering researchers a powerful tool for hypothesis generation and scientific discovery.

## 1. Introduction

A key open question in neuroscience is whether cognitive processes can be inferred from brain activation patterns— a problem known as reverse inference (Poldrack, 2006). Large-scale meta-analysis has facilitated reverse inference through meta-analytic functional decoding, a method that offers a quantitative estimation of behavioral profiles associated with brain regions (Amft et al., 2015; Bzdok et al., 2013b, 2013a; Cieslik et al., 2013; Laird et al., 2009; Nickl-Jockschat et al., 2015; Peraza et al., 2024; Poldrack, 2011; Rottschy et al., 2013; Smith et al., 2009). However, existing functional decoding methods are fundamentally limited by the challenge of how to model unstructured text from publications to capture discrete mental states that align with established theoretical frameworks.

Coordinate-based meta-analytic (CBMA) databases, such as BrainMap (Fox et al., 2005; Laird et al., 2011, 2009, 2005) and Neurosynth (Yarkoni et al., 2011), along with the Cognitive Atlas ontology (Poldrack et al., 2011), provide robust data to facilitate quantitative meta-analytic functional decoding in neuroimaging, while recent overviews of neuroimaging meta-analysis have highlighted both the continuing value of large-scale synthesis and the need for improved coverage, quality control, and integration across coordinate- and image-based resources (Oudyk et al., 2025). The implementation of meta-analytic functional decoding methods on popular websites (e.g., Neurosynth (Yarkoni et al., 2011) and NeuroVault (Gorgolewski et al., 2015)), as well as meta-analysis software (e.g., NiMARE (Salo et al., 2023)), has boosted their popularity within the neuroimaging community.

However, current methods present several limitations. For instance, the most commonly used methods, correlation-type decoders, are not based on a formal predictive model, do not explicitly learn latent structures associated with cognitive processes, and are constrained by precomputed meta-analytic maps and fixed vocabularies rather than learned representations optimized for prediction. The Generalized Correspondence Latent Dirichlet Allocation (GC-LDA) approach (Rubin et al., 2016, 2017) addresses some of these limitations by learning joint probabilities from latent variables in the model, allowing for predictions on unseen brain patterns. Nevertheless, GC-LDA is significantly influenced by the data, context, and assumptions established before model training, such as spatial priors, the number of topics, and other model parameters. Additionally, GC-LDA operates entirely unsupervised, suggesting that the model is not optimized for classification or prediction accuracy. Thus, other models refined to enhance prediction parameters should outperform GC-LDA (Rubin et al., 2017).

In addition, the effectiveness of functional decoding is held back by its reliance on outdated Natural Language Processing (NLP) techniques. For instance, it is quite common for text extracted from neuroimaging publications to be analyzed using term-frequency inverse document frequency (TF-IDF) (Aizawa, 2003), a classic metric that measures a word’s importance based on its frequency. While TF-IDF has proven helpful in large-scale meta-analysis (Dockès et al., 2020; Rubin et al., 2016; Salo et al., 2023; Yarkoni et al., 2011), as a “*bag of words*” method, it provides only a sparse view of word representation, fails to capture semantics, and overlooks relationships between words within the same conceptual family. Many widely used functional decoding approaches, such as the Neurosynth correlation decoder and GC-LDA, are built on similar bag-of-words representations derived from term frequencies, and therefore inherit these limitations. Additionally, its vocabulary is confined to words present in the neuroscience corpora, restricting the inclusion of external terms and, more importantly, preventing the use of richer representations such as phrases, sentences, and term definitions.

The emergence of large language models (LLMs) (Liu et al., 2023; Vaswani et al., 2017) has transformed the field of NLP and is now poised to revolutionize neuroimaging meta-analysis. By capturing deep semantic relationships, LLMs overcome the limitations of traditional NLP metrics like TF-IDF, providing rich contextual embeddings that understand relationships between related concepts and capture richer semantics of phrases and definitions (Gunasekar et al., 2023; Srivastava et al., 2023). LLMs have been applied to neuroimaging meta-analysis to enhance text encoding and synthesize results from various articles (Ngo et al., 2021). In neuroscience, BrainGPT, an LLM fine-tuned on neuroscience literature, has demonstrated domain-specific advantages by outperforming experts in predicting neuroscience findings (Luo et al., 2024).

Recently, NeuroConText (Ghayem et al., 2026) introduced a CLIP-based (Radford et al., 2021) framework for text-to-brain association that significantly outperformed previous approaches including NeuroQuery (Dockès et al., 2020) and Text2Brain (Ngo et al., 2021). By using a pretrained LLM transformer to encode text, coordinate-derived modeled activation maps to represent brain activity, and brain parcellation for dimensionality reduction, NeuroConText demonstrated improved flexibility in handling longer texts and better text-brain associations. Importantly, NeuroConText established that contrastive learning can align unstructured neuroimaging text with brain activation patterns in a shared latent space. However, its primary focus was text-to-brain association, and it did not systematically evaluate brain-to-text decoding or examine whether domain-specific LLMs such as BrainGPT could further optimize the underlying text–brain representation.

In parallel, ontology-based decoding approaches have shown that structured cognitive vocabularies can provide a principled framework for reverse inference. Varoquaux et al. (2018) introduced an ontology-guided multi-label decoding framework for building atlases of cognition, and Menuet et al. (2022) later combined NeuroVault statistical maps with Cognitive Atlas annotations to decode mental processes in unseen studies, including the use of a reduced Cognitive Atlas ontology. These studies established important foundations for functional decoding by showing the utility of structured ontological cognitive labels for interpretable associations between brain activity and mental processes. However, these ontology-based frameworks have primarily been developed in image-based meta-analysis settings, which, although powerful due to their use of full statistical maps, remain limited in breadth by the small proportion of published studies that share statistical images in open repositories such as NeuroVault. Conversely, NeuroConText leverages the broader coordinate-based literature through contrastive text–brain learning, but does not incorporate an ontology-guided decoding framework for mapping brain activation patterns onto structured cognitive tasks, concepts, and domains.

In this work, we present the NiCLIP (Neuroimaging Contrastive Language-Image Pre-training) model, an ontology-guided CLIP-based framework for functional decoding of brain activation maps. NiCLIP combines contrastive learning with an ontology-guided probabilistic decoding framework for predicting cognitive tasks, concepts, and domains from brain activation patterns. Specifically, NiCLIP builds on the contrastive text–brain representation strategy introduced by NeuroConText, aligning text from neuroimaging articles with coordinate-derived brain activation maps in a shared latent space, and extends this backbone to support structured reverse inference using a standardized cognitive vocabulary. The central contribution of NiCLIP is therefore the integration of contrastive text–brain alignment with ontology-guided decoding, allowing brain activation maps to be interpreted at multiple levels of cognitive abstraction. We evaluated NiCLIP using more than 23,000 neuroimaging articles with full text and reported activation coordinates from PubMed Central. First, we assessed how choices in the underlying representation model, including article section and LLM type, affected text–brain alignment, treating domain-specific LLMs such as BrainGPT as a potential optimization of the backbone. Second, we assessed the external validity of NiCLIP by evaluating its functional decoding performance on independent task-fMRI group-average maps from the Human Connectome Project (HCP) (Barch et al., 2013; Smith et al., 2013; Uğurbil et al., 2013; Van Essen et al., 2013, 2012) using a vocabulary derived from the Cognitive Atlas ontology (Poldrack et al., 2011). In this evaluation, we examined how LLM choice, article section used for model training (i.e., abstract vs. body), and cognitive ontology affected the decoder’s predictions of cognitive tasks, concepts, and domains. We then conducted additional analyses to characterize NiCLIP’s performance across different types of input brain maps, including six anatomical regions defined by meta-analytic parcellations and subject-level HCP activation maps. Finally, we compared NiCLIP against established baseline models, including the Neurosynth correlation decoder and the GC-LDA decoder.

## 2. Results

### 2.1. Overview

**Fig. 1** provides an overview of our general framework to train the text-to-brain model (**Fig. 1A**) and perform functional decoding of a brain activation map (**Fig. 1B**). To train the CLIP model, we required text and image data. First, we searched the PubMed Central open-source collections for fMRI articles. Second, we extracted the text and activation coordinates from the downloaded articles using Pubget (Dockès et al., 2024). Third, we identified text and brain activation embeddings for each article, along with corresponding coordinates. For the text features, we used a pre-trained LLM. Meanwhile, the brain activation features were obtained by transforming the coordinates into a modeled activation map using MKDA (Wager et al., 2007) methods and reducing the image dimensionality with a continuous brain parcellation defined by the DiFuMo 512 regions atlas (Dadi et al., 2020). Finally, we entered the normalized text and brain embeddings into the CLIP model to learn a shared latent space for the text and image encoders.

**Figure 1.**
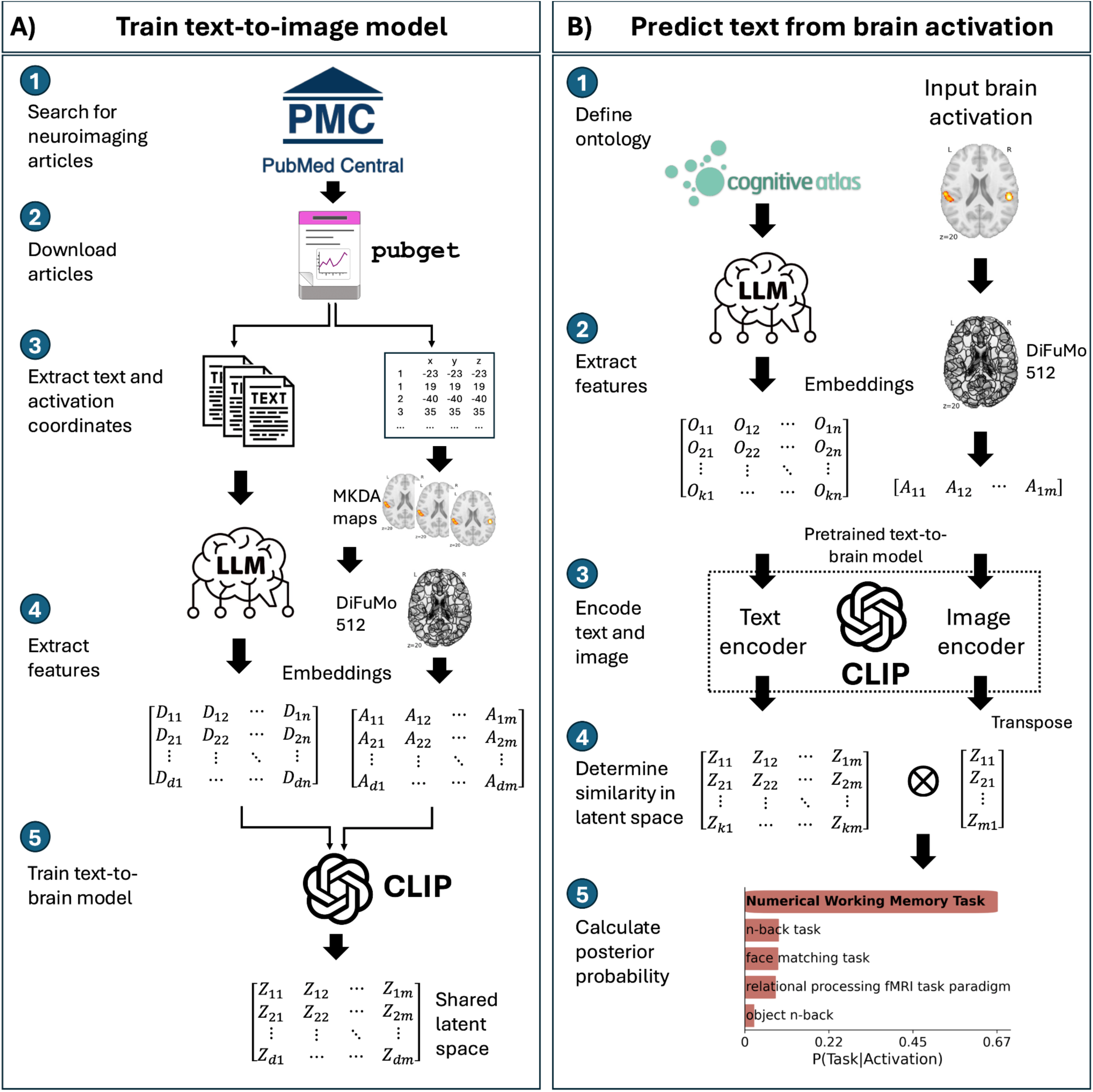
Overview of the framework for training the text-to-brain model and decoding brain activation maps. **(A)** The text-to-brain CLIP model was trained using text and brain activation coordinates sourced from a collection of fMRI articles downloaded from PubMed Central. Pubget was employed to download and preprocess the articles in a standardized format. Text embeddings were determined using a pre-trained LLM. Image embeddings were obtained by first calculating an MKDA-modeled activation brain map, and second applying a continuous brain parcellation defined by the DiFuMo 512 atlas. **(B)** The brain decoding model relies on a cognitive ontology to predict text from input brain activation. The embeddings of task names along with their definitions were extracted using an LLM transformer, while image features were determined using the DiFuMo brain parcellation. The text and image embeddings were processed through the pre-trained text and image encoders from CLIP, yielding new embeddings in a shared latent space. The posterior probability representing the predicted task from the input activation was calculated based on the similarity between the text and image embeddings in the shared latent space.

To make predictions based on brain activation, we needed a cognitive ontology that consisted of fMRI tasks with their respective definitions, along with concepts associated with the tasks and their corresponding high-level domains. We first obtained a joint embedding of the task name and definition using a pre-trained LLM. The input brain activation embedding was determined using the DiFuMo 512 atlas. Then, the cognitive ontology and input activation embeddings were passed through the text and image encoders from the pre-trained CLIP model. Next, we determined the cosine similarity between the text and image embeddings in the shared latent space. This similarity was used to calculate the posterior probability of a task given an activation, given the shared latent space.

### 2.2. CLIP model evaluation for text-to-brain association

To evaluate the quality of the learned text–brain alignment of the underlying trained CLIP we used two complementary retrieval-based metrics: Recall@k and Mix&Match (Ghayem et al., 2026). Recall@k measures the proportion of cases in which the correct brain activation map is retrieved among the top k most similar candidates given a text query, reflecting the ranking performance of the model (e.g., Recall@10 and Recall@100 correspond to retrieval within the top 10 and 100 candidates, respectively). In contrast, Mix&Match evaluates the model’s ability to correctly associate multiple text and brain activation pairs simultaneously. In this setting, a batch of text and brain embeddings is randomly shuffled, and the model must recover the correct one-to-one correspondences based on similarity in the shared latent space. This metric captures the global consistency of the learned alignment beyond top-k retrieval. Higher values for both metrics indicate better alignment between text and brain representations. Models were evaluated using a cross-validation framework on a dataset of 23,865 articles (see Methods for details).

Four different large language models (i.e., BrainGPT-7B-v0.2, Mistral-7B-v0.1, BrainGPT-7B-v0.1, and Llama-2-7b-chat-hf) were evaluated for text features extraction across two document sections (i.e., abstract and body). Training the CLIP model with the body portion of the articles and embeddings extracted with BrainGPT-7B-v0.2 demonstrated the strongest performance across all metrics (Recall@10: 33.56±4.12, Recall@100: 71.95±4.18, Mix&Match: 90.04±1.66), closely followed by its foundational LLM, Mistral-7B-v0.1 (Recall@10: 33.36±4.05, Recall@100: 71.78±4.22, Mix&Match: 89.97±1.70). BrainGPT-7B-v0.1 and Llama-2-7b-chat-hf showed comparable but slightly lower performance, with Llama-2-7b-chat-hf marginally outperforming BrainGPT-7B-v0.1 on Recall@k and Mix&Match. The Mistral-7B-v0.1 condition represents the closest NeuroConText-equivalent setting in our evaluation, as it uses the same base LLM and the same general CLIP-style text–brain alignment strategy; thus, this comparison assesses whether alternative text encoders improve the shared representation backbone relative to the NeuroConText-style configuration.

When using only the abstract sections to train the CLIP model, all LLMs demonstrated lower performance than when training with the full body of the articles. BrainGPT-7B-v0.2 again achieved the highest scores (Recall@10: 24.01±1.84, Recall@100: 61.98±4.28, Mix&Match: 85.82±1.84), with BrainGPT-7B-v0.1 performing slightly better than Mistral and Llama models across all three metrics. Llama-2-7b-chat-hf consistently showed the lowest performance in the abstract section evaluation.

Across all LLM configurations, performance differences were small (**Table 1**). BrainGPT-7B-v0.2 showed numerically higher text-to-brain association scores compared to the other LLMs; however, the differences were within the range of cross-validation variability and no strong conclusions can be drawn about the superiority of any single LLM based on these results alone. The larger and more consistent effect was observed for the article section: models trained using the full text body outperformed those trained using abstracts alone across all LLMs. Distributions of the evaluation metrics presented in **Table 1** are available in the supplementary information (**Fig. S1**).

**Table 1.** Text-to-brain association performance of CLIP models across LLMs and article sections. The CLIP backbone was evaluated using text embeddings derived from either the full article body or the abstract, and from four LLMs: BrainGPT-7B-v0.2, Mistral-7B-v0.1, BrainGPT-7B-v0.1, and Llama-2-7b-chat-hf. Recall@10 and Recall@100 measure the percentage of cases in which the correct brain activation map was retrieved among the top 10 or top 100 candidates given the article text embedding. Mix&Match measures whether the model correctly recovers text–brain correspondences after candidate pairs are shuffled, reflecting the global consistency of the learned shared latent space. Values represent mean ± standard deviation across cross-validation sets. Higher values indicate better text–brain alignment. The best-performing value for each metric is highlighted in bold.

| CLIP model setting |  | Metrics |  |  |
| --- | --- | --- | --- | --- |
| Section | LLM | Recall@10 | Recall@100 | Mix&Match |
| Body | BrainGPT-7B-v0.2 | **33.56 ± 4.12** | **71.95 ± 4.18** | **90.04 ± 1.66** |
|  | Mistral-7B-v0.1 | 33.36 ± 4.05 | 71.78 ± 4.22 | 89.97 ± 1.70 |
|  | BrainGPT-7B-v0.1 | 32.30 ± 4.16 | 70.66 ± 4.81 | 89.65 ± 1.83 |
|  | Llama-2-7b-chat-hf | 32.50 ± 4.08 | 71.08 ± 4.81 | 89.72 ± 1.84 |
| Abstract | BrainGPT-7B-v0.2 | 24.01 ± 1.84 | 61.98 ± 4.28 | 85.82 ± 1.84 |
|  | Mistral-7B-v0.1 | 23.36 ± 2.12 | 61.37 ± 4.27 | 85.57 ± 1.88 |
|  | BrainGPT-7B-v0.1 | 23.51 ± 1.98 | 61.94 ± 4.09 | 85.73 ± 1.79 |
|  | Llama-2-7b-chat-hf | 22.46 ± 2.07 | 60.69 ± 4.13 | 85.32 ± 1.87 |

**Note:** Values represent mean ± standard deviation across cross-validation sets. The best-performing value for each metric is highlighted in bold font.

### 2.3. NiCLIP model settings evaluation

Building on the CLIP model evaluation described above, we next assessed the performance of NiCLIP, which extends the learned text–brain alignment to enable reverse inference (brain-to-text decoding) through a structured cognitive ontology. Here, we incorporated the Cognitive Atlas to define tasks, concepts, and domains, where a “domain” represents a high-level cognitive category. The ontology includes fMRI tasks and their definitions, along with associated concepts and higher-level domains. Task predictions are based on the semantic representation of task names and definitions, while concept and domain predictions are derived from task-to-concept and concept-to-domain mappings.

We evaluated the NiCLIP framework across different parameter configurations to assess decoding performance across seven cognitive domains from the HCP task fMRI dataset (i.e., emotion, gambling, language, motor, relational, social, and working memory), and compared results to two baseline models (Neurosynth and GC-LDA). First, we evaluated CLIP model configurations identified in the previous analysis (**Table 1**), including article section (abstract vs. body) and choice of pre-trained LLM (BrainGPT-7B-v0.2, Mistral-7B-v0.1, BrainGPT-7B-v0.1, and Llama-2-7b-chat-hf). Second, we compared two versions of the Cognitive Atlas ontology: the full ontology and a reduced version containing the most commonly used fMRI tasks with more robust concept-to-task mappings. Third, we assessed two text embedding strategies: using the task name alone versus the task name combined with its definition. Results are summarized in **Table 2**.

**Table 2.**
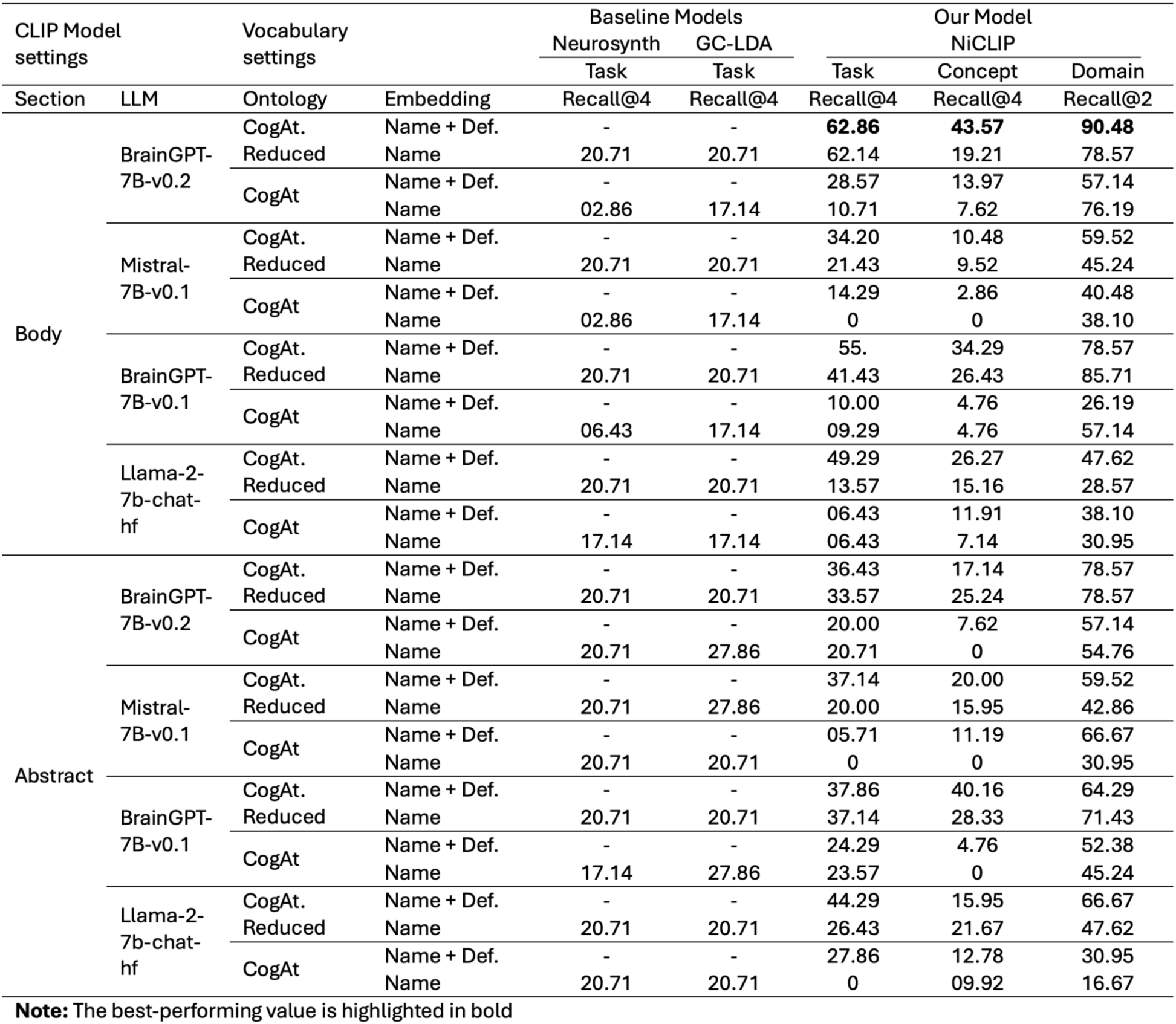
Functional decoding performance was evaluated on seven HCP task-fMRI domains: emotion, gambling, language, motor, relational, social, and working memory. Rows show NiCLIP configurations varying by article section used for CLIP training, LLM text encoder, ontology version, and task embedding strategy. “CogAt” refers to the original Cognitive Atlas ontology, whereas “CogAt. Reduced” refers to the curated reduced ontology with more complete task-to-concept mappings. “Name + Def.” indicates that task embeddings were computed from both the task name and definition, whereas “Name” uses the task name alone. Baseline models include the Neurosynth correlation decoder and GC-LDA, which provide task-level predictions only. NiCLIP performance is reported for task, concept, and domain predictions. Task and concept decoding are evaluated with Recall@4, and domain decoding is evaluated with Recall@2. Higher values indicate better decoding performance. Dashes indicate configurations that were not applicable to the corresponding baseline model. The best-performing value for each NiCLIP prediction level is highlighted in bold.

The highest performance across all categories was achieved using the complete text of the articles to train CLIP, BrainGPT-7B-v0.2, combined with the reduced Cognitive Atlas ontology, along with their definitions and task names (Task: 62.86%, Concept: 43.57%, Domain: 90.48% at Recall@2), substantially outperforming both baseline models with over a 40% improvement in recall. BrainGPT-7B-v0.1 achieved the second-best performance (Task: 55%, Concept: 34.29%, Process: 78.57%), indicating consistent gains from domain-specific LLMs.

Across all models, the curated version of Cognitive Atlas, which retains the most commonly used fMRI tasks while enriching their concept mappings, consistently outperformed the original Cognitive Atlas ontology across all models, with additional gains when using both the task name and definition rather than the name alone. In contrast, some configurations with the full Cognitive Atlas ontology exhibited minimal or zero recall, particularly with Mistral-7B-v0.1. Baseline models (Neurosynth and GC-LDA) demonstrated limited effectiveness, with maximum Recall@4 scores of 20.71% for tasks–substantially lower than NiCLIP. However, note that because **Table 2** reports all NiCLIP ablation settings, some suboptimal NiCLIP configurations performed worse than baseline decoders.

Models trained using only the abstract, showed generally lower performance compared to those trained on the full-text body, although similar patterns were observed for both. BrainGPT-7B-v0.1 with CogAt reduced ontology and “name + definition” vocabulary achieved the highest concept recall (40.16%), while Llama-2-7b-chat-hf excelled in task predictions recall (44.29%) with the same configuration. The predictions of domains consistently showed higher recall rates than tasks and concepts across all models and configurations, with BrainGPT-7B-v0.2 also achieving the best performance (78.57%) when trained with the abstract section, utilizing the reduced ontology with both task name only and name plus definition. However, note that raw Recall@K values are not directly comparable across tasks, concepts, and domains because these vocabularies differ substantially in size. Domain prediction is structurally easier because there are fewer candidate domain labels, so higher domain recall reflects both model performance and the smaller label space.

### 2.4. Evaluating NiCLIP for predicting text from different brain maps

#### 2.4.1. Group-level activation maps

Diving deeper into the previous findings, **Fig. 2** shows the top 5 predictions of NiCLIP, sorted by the magnitude of the posterior probabilities for task (P(T|A)), concept (P(C|A)), and domain (P(D|A)). The probability of P(T|A) is defined by Bayes’ theorem as the normalized joint probability of P(A|T) and the prior P(T). The likelihood P(A|T) reflects the similarity in the latent space of the activation embedding to the embeddings of task names and definitions from the ontology. At the same time, the prior distribution indicates the prevalence of the tasks in the literature. The probabilities for concepts and domains are not based on their semantic content, like the task itself, but instead on the probability of the task and their connection established in the ontology using the noisy-OR model (additional details are presented in the Methods. For these results, we used the best-performing CLIP model from the previous analysis. Additional comparisons between the NiCLIP task predictions and baseline models are available in the supplementary information (**Fig. S2**).

**Figure 2.**
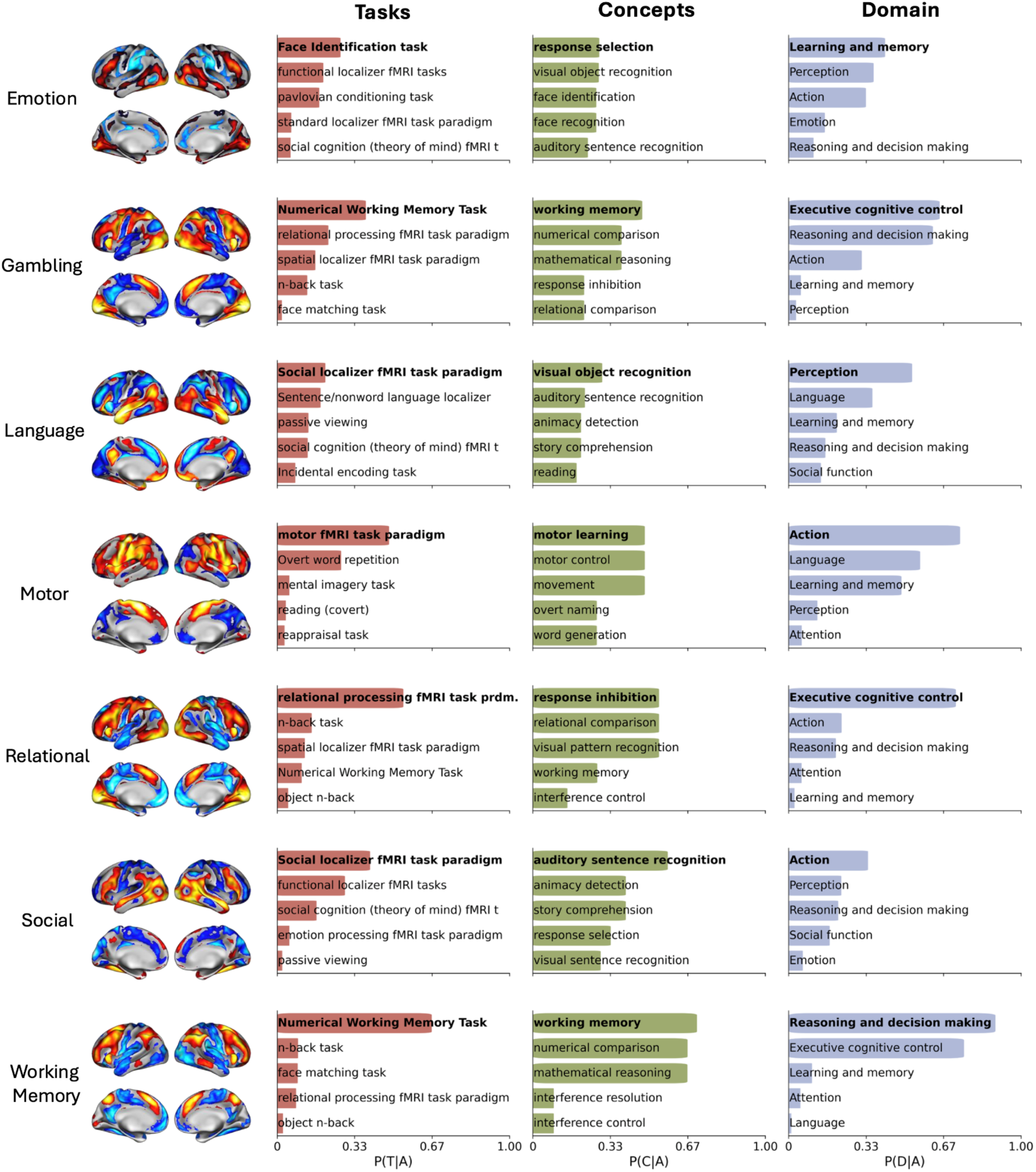
NiCLIP predicts tasks, concepts, and domains from brain activation patterns based on group-level maps. This analysis provides prediction probabilities across seven major cognitive domains from the Human Connectome Project (HCP) task fMRI dataset. For each domain (Emotion, Gambling, Language, Motor, Relational, Social, and Working Memory), we present three types of predictions: the probability of a task given an activation pattern (P(T|A)), the probability of a concept given an activation (P(C|A)), and the probability of a domain given an activation (P(D|A)). Each prediction is illustrated with horizontal bars representing prediction strength, showcasing the top five predictions for each category. Task labels correspond to Cognitive Atlas task names displayed for readability; the semantic embeddings used for decoding incorporate both task names and task definitions.

We decoded seven HCP task contrasts to evaluate NiCLIP’s predictions (**Fig. 2**). For the emotion processing task (“Face vs. Shape”), NiCLIP identified face identification (27.4%) as the top task, followed by a functional localizer task (19.9%). Related concepts included response selection (28.6%), visual object recognition (28.5%), and face recognition (27.4%), which map to Learning and Memory (41.6%) and Perception (36.7%) domains. The gambling task (“Reward vs. Baseline”) was mainly associated with numerical working memory tasks (38.2%) and relational processing (22.1%). Key concepts involved working memory (47.0%), numerical comparison (38.3%), and mathematical reasoning (38.2%), with Executive Cognitive Control (65.0%) and Reasoning and Decision-Making (62.2%) as the top domains. For language (“Story vs. Math”), NiCLIP predicted social localizer (18.9%) and language localizer task (17.0%), with concepts including visual object recognition (27.4%), auditory sentence recognition (24.6%), and reading (20.5%). The motor task showed expected links with motor fMRI task paradigm (48.3%) and motor-related concepts (movement, motor control, motor learning; all 48.3%), mapping to Action (73.8%) and Language (56.6%) domains. The relational processing task identified the relational processing fMRI task paradigm (54.5%) with Executive Cognitive Control (72.0%) as the primary domain. Social cognition (“TOM vs. Random”) predicted social localizer (39.9%) with concepts including auditory sentence recognition (58.0%) and animacy detection (39.9%). Finally, working memory (“2-Back vs. 0-Back”) showed strong associations with numerical working memory tasks (67.2%) and working memory concepts (71.0%), mapping to Reasoning and Decision-Making (89.5%) and Executive Cognitive Control (75.8%).

#### 2.4.2. Region of interest maps

Next, we present a potential application of reverse inference methods for exploring novel hypotheses and discovering tasks, concepts, and domains associated with a specific region of interest (ROI). For this experiment, we also utilized the best-performing CLIP model (i.e., BrainGPT-7B-v0.2), which was trained on the complete body of articles. The NiCLIP model relied on a reduced version of the Cognitive Atlas as its cognitive ontology. NiCLIP predictions on ROIs were tested on six popular anatomical brain regions, including the amygdala (Bzdok et al., 2013a), hippocampus (Plachti et al., 2019), insula (Chang et al., 2013), striatum (Liu et al., 2020), right temporoparietal junction (rTPJ) (Bzdok et al., 2013b), and ventromedial prefrontal cortex (vmPFC) (Chase et al., 2020). **Fig. 3** presents the results of predicting tasks, concepts, and domains across the six ROIs. Unlike the previous predictions on dense maps from **Fig. 2**, the top predicted task here resulted in a more distinct probability relative to the second-highest task. Additional comparisons between the NiCLIP task predictions and the baseline models are provided in the supplementary information (**Fig. S3**).

**Figure 3.**
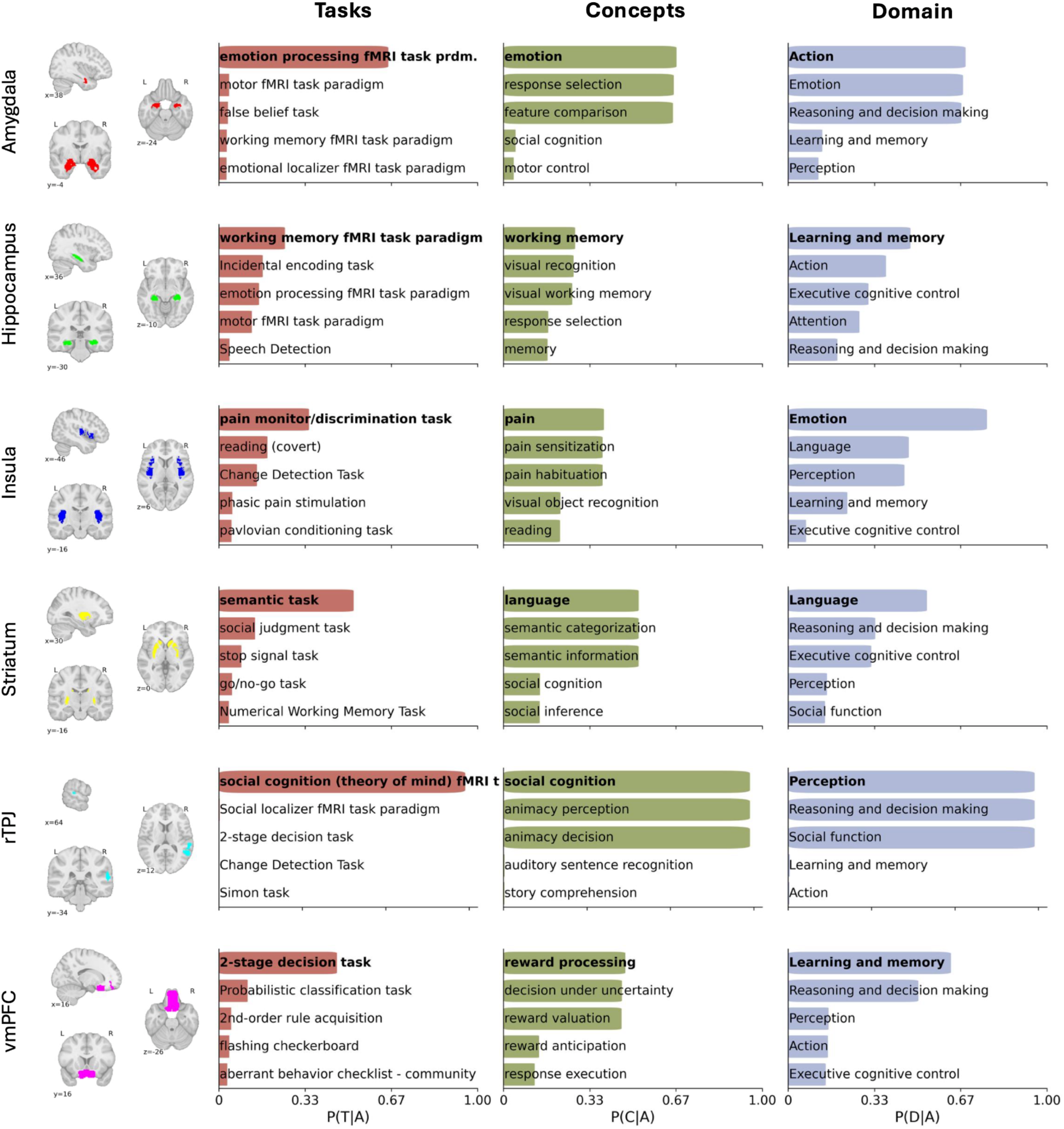
NiCLIP predicts tasks, concepts, and domains from brain ROIs. We conducted a comprehensive analysis of prediction probabilities across six different ROIs. For each ROI (amygdala, hippocampus, insula, striatum, rTPJ, vmPFC), we display three types of predictions: the probability of a task given an activation pattern (P(T|A)), the probability of a concept given an activation (P(C|A)), and the probability of a domain given an activation (P(D|A)). Each prediction is visualized with horizontal bars indicating prediction strength, with the top five predictions shown for each category. Task labels correspond to Cognitive Atlas task names displayed for readability; the semantic embeddings used for decoding incorporate both task names and task definitions.

We analyzed NiCLIP predictions across six anatomical ROIs to demonstrate its ability to identify regional functional specialization (**Fig. 3**). The amygdala showed a strong association with emotion processing tasks (65.5%), with other tasks below 5%. Top concepts included emotion, response selection, and feature comparison (∼65%), aligning with Action (68.6%), Emotion (67.6%), and Reasoning and Decision-Making (67.0%) domains. The hippocampus was linked to working memory fMRI task (25.6%), incidental encoding (17.0%), emotion processing (15.6%), and motor tasks (12.7%), with memory-related concepts (working memory, visual recognition, memory) dominating. Learning and Memory was the primary domain (47.2%), followed by Action and Executive cognitive control (31.0%). The insula exhibited pain-related functions, including pain monitoring/discrimination (34.9%) and related concepts (pain, pain sensation, pain habituation; ∼38% each), with a strong mapping to the Emotion domain (76.9%). The striatum showed language specialization, connecting with semantic tasks (52.1%) and language-related concepts (52.2%), with Language (53.6%) as the dominant domain. The rTPJ displayed remarkable selectivity for social cognition/theory of mind tasks (98.5%), with related concepts (social cognition, animacy perception) mapping to Perception, Reasoning and Decision-Making, and Social Function domains (all 98.5%). Finally, the vmPFC was associated with decision-making tasks (2-stage decision: 45.7%), reward-related concepts (reward processing: 47.2%, decision-making under uncertainty: 45.9%), and the domains of Learning and Memory (63%) and Reasoning and Decision-Making (50.4%).

#### 2.4.3. Subject-level activation maps

Following the prediction of group-level activation maps, we explored whether NiCLIP can identify the underlying tasks, concepts, and domains in subject-level images. In practice, researchers typically report activation foci from group-level images of significant findings in scientific papers. Consequently, our model is capable of producing robust predictions for population-level average activation maps, as illustrated in **Fig. 2**. In contrast, subject-level images exhibit high variability and noise, and individual subject differences are known to result in distinct brain activation patterns within the same task and condition. We anticipated that these factors could potentially interfere with NiCLIP’s predictions. For this experiment, we also utilized the best-performing CLIP model. We conducted a qualitative evaluation of the NiCLIP predictions on the exact seven major domains from the HCP tasks fMRI dataset, but using the subject-level activation maps instead. These domains included tasks along with their respective contrasts of interest, including emotion, gambling, language, motor, relational, social, and working memory. In this analysis, we included 787 subjects and the distribution of the Recall@K score is presented in **Fig. S4**. **Fig. S5** illustrates the results for one particular subject with the best performance and shows the top 5 predictions sorted by the magnitude of the posterior probabilities (i.e., P(T|A), P(C|A), and P(D|A)). Overall, functional decoding on subject-level brain activation maps was less effective than the results on group-level images, with an average Recall@4 score of 38.19% for tasks, a Recall@4 score of 25.34% for concepts, and a Recall@2 score of 52.01% for the domain. Correct findings were rarely ranked among the top five predictions for most of the target brain images (**Fig. S5**). Additional comparisons between the NiCLIP task predictions and the baseline models (**Fig. S6**), as well as visualizations of the embeddings geometry (**Figs. S7 and S8**), are provided in the supplementary information.

#### 2.4.4. Different variations of the same activation map

Finally, we examined the extent of NiCLIP’s capability in predicting subject-level activation maps. To achieve this, we utilized the group-level activation task from the motor domain of the HCP dataset and created six distinct variations of the original map. One map included only the positively activated region, while another included only the negatively activated region. We also tested a map with the positively activated regions of the original maps, but with the sign flipped. Lastly, we evaluated the prediction on three different ROI maps, representing the top 10, 5, and 1 percentiles of the positively activated regions. **Fig. S9** shows the results of the top 5 predictions sorted by the magnitude of the posterior probabilities (i.e., P(T|A), P(C|A), and P(D|A)). Overall, the six variations produced different results. Decoding the negative tail alone, or the positive tail with the sign flipped, did not produce a motor task in the top 5 predictions. The ROI created by taking the top 10 and top 5 percentiles included the “motor fMRI task paradigm” in the top three, but the top two predicted tasks did not match the original image’s prediction. The map consisting of the positive tail alone was closer to the original image’s prediction. The ROI comprising the top one percentile of positive activation was the only variation that showed the exact prediction and selectivity as the original image, differing only in the order of the top two tasks.

## 3. Discussion

The topic of formal reverse inference on brain activation maps has remained an open question in the neuroscience community for many years. Although several attempts have been made to address this gap, mostly utilizing emerging meta-analytic approaches and databases, multiple limitations persist in meta-analytic methods for consolidating existing data for functional decoding. Additionally, the absence of a reliable cognitive ontology has hindered broader adoption and application. To tackle these current challenges, we present NiCLIP in this work, a contrastive language-image pretraining model designed to predict fMRI tasks, concepts, and domains from brain images.

We compared four different LLM models and found that domain-specific fine-tuned LLMs (BrainGPT models) and their general-purpose base models (Mistral-7B-v0.1) showed numerically similar performance within the NiCLIP framework, consistent with the observation that modern open-source LLMs already encode rich semantic structure relevant to neuroimaging. The primary contribution of NiCLIP lies instead in the ontology-guided decoding framework: CLIP-derived similarities are converted into task posterior probabilities and then propagated through task-to-concept and concept-to-domain mappings to produce structured functional decoding outputs. We evaluated the predictions of NiCLIP models on group-level activation maps from the HCP and found overall accurate predictions of tasks, concepts, and domains.

However, when NiCLIP was applied to the same tasks on subject-level images, it produced inconsistent results, with accurate task predictions only for a few tasks, such as emotion and social, while yielding incorrect predictions for others. Finally, we investigated the capabilities of the NiCLIP model for decoding regions of interest. In that experiment, we successfully predicted tasks, concepts, and domains across six different meta-analytic ROIs tested. An extensive evaluation of NiCLIP revealed that it is crucial not only to utilize a fine-tuned LLM and train the CLIP model on the full text of the articles, but also to leverage a curated cognitive ontology with precise task-to-concept and concept-to-domain mappings. Taken together, our findings situate NiCLIP as a flexible ontology-guided framework for functional decoding that extends contrastive text–brain alignment beyond retrieval and toward structured reverse inference over tasks, concepts, and domains.

### 3.1. Situating NiCLIP within current tools for decoding brain images

NiCLIP advances current functional decoding approaches by combining contrastive text–brain alignment with ontology-guided probabilistic decoding. Existing functional decoding methods, such as correlation-type decoders, are constrained by precomputed meta-analytic maps and fixed vocabularies rather than learned representations optimized for prediction. Other methods, like GC-LDA, operate entirely unsupervised, indicating that these models are not optimized for classification or prediction accuracy. Generally, these methods are limited by the small number of pre-created meta-analytic maps used for predictions, rely on TF-IDF to define concepts or maps—which ignores semantic context—and operate within a limited vocabulary, thereby preventing the prediction of external or unseen concepts. These limitations reflect the broader constraints of bag-of-words text representations. In contrast, replacing TF-IDF with LLM-derived contextual embeddings leads to more accurate and semantically meaningful functional decoding, with the best NiCLIP configuration achieving more than three times the task-level recall of both Neurosynth and GC-LDA baselines (Table 2), demonstrating that the CLIP-based approach provides substantial empirical advantages over simpler term-frequency methods. As a CLIP-based decoding model, NiCLIP utilizes an image-text latent space to make predictions on unseen data. CLIP models are also self-supervised, meaning their parameters are optimized to match text to corresponding images with minimal training error and they are fine-tuned for accurate text-image matching on unseen data. NiCLIP models are not constrained by a set of pre-computed meta-analytic maps since the model operates by finding similarities within the shared latent space. Both LLM and CLIP models enable linking the entire abstract and text body of a publication to brain activation patterns, thereby overcoming the limitations of bag-of-words and TF-IDF methods. Additionally, LLM enables the use of external terms and their definitions to annotate publication texts, for example, concepts from established ontologies such as the Cognitive Atlas.

Although the NeuroQuery framework offers a solution to expand to external terms based on similarity (Dockès et al., 2020), such expansions remain limited to simple term combinations and do not fully capture entire ontologies or phrases. Additionally, the similarities between terms in NeuroQuery are only based on their names. In contrast, NiCLIP utilizes both the term names and their definitions, providing more precise and distinct semantic content. By employing a comprehensive ontology and leveraging semantic similarities between tasks and their descriptions from a cognitive ontology, we can also expand predictions to include concepts and domain-specific distributions, thereby providing additional context. NiCLIP also introduces a predictive framework for identifying associations with tasks that lack sufficient meta-analytic data to appear in traditional decoding results. For example, the n-back task was among the top predictions made by NiCLIP for the working memory activation maps. However, the “n-back task” was underrepresented in the literature; as a result, no meta-analytic maps were available for the Neurosynth correlation decoder to establish associations with the working memory maps, as shown in the supplementary information (**Fig. S2**).

While recent decoding approaches using NeuroVault statistical maps and Cognitive Atlas have shown improved performance over traditional CBMA-based methods (e.g., Neurosynth and GC-LDA) (Mensch et al., 2021, 2017; Menuet et al., 2022; Varoquaux et al., 2018), they face significant limitations that restrict their usefulness for comprehensive functional decoding. Despite efforts within the community, most neuroimaging studies do not share their statistical maps, resulting in NeuroVault containing only a limited set of tasks and cognitive domains with sparse coverage (Salo et al., 2023). Moreover, existing NeuroVault data suffers from inconsistent metadata annotation and frequent mislabeling of image modalities, map types, and Cognitive Atlas tasks (Peraza et al., 2025). These quality and coverage issues fundamentally restrict the generalizability of NeuroVault-based decoders. In contrast, our CBMA-based approach benefits from the extensive coverage of coordinate-based databases, which currently comprise the largest collection of neuroscience publications covering diverse fMRI tasks and cognitive domains (Oudyk et al., 2025). This broad literature coverage enables NiCLIP to decode activation patterns across a wider range of cognitive functions, including those underrepresented in NeuroVault. While whole-brain statistical maps offer richer spatial information, the balance between data richness and domain coverage makes CBMA more suitable for applications that require diverse functional decoding. Therefore, rather than a limitation, our CBMA foundation offers a strategic advantage for uncovering associations across the full spectrum of cognitive neuroscience, reducing the selection bias associated with repository-dependent decoding approaches.

### 3.2. The need for better ontologies

One key feature of the current framework is its ability to make predictions using external terminology, such as terms provided by an ontology. Here, NiCLIP uses a cognitive ontology to define the fMRI tasks and their connections to concepts and domains. Despite the extensive collection of tasks and concepts, the Cognitive Atlas database remains limited. As a community-driven ontology, these mappings reflect the opinions of individual researchers and are not necessarily factual (Poldrack et al., 2011). We might find popular tasks, such as the “motor fMRI task paradigm”, linked only to working memory concepts, while missing relevant terms like movement and motor, among others. The task names and descriptions are still not standardized. Some tasks may have definitions made up of two sentences, while others consist of multiple sentences, which affects the final embedding computed for prediction in the shared latent space.

We demonstrated that a reduced and curated representation of the Cognitive Atlas tasks, combined with a more robust and comprehensive mapping of concepts, outperforms the original Cognitive Atlas ontology. This highlights the importance of cognitive ontology for reverse inference tasks. Following the trend of leveraging LLMs for such challenging tasks, we believe that a data-driven ontology derived from the entire neuroscience literature has the potential to improve existing cognitive ontologies by providing more accurate and standardized task names and definitions, along with better task-to-concept mappings. Beyond improving these mappings, an alternative is to bypass the task-to-concept propagation layer altogether, computing cosine similarities directly between brain embeddings and concept-level text embeddings; this could reduce the impact of imperfect task-to-concept mappings and may yield more accurate concept-level predictions. Either way, developing more standardized and precise cognitive ontologies remains crucial to fully harness the potential of LLM-powered functional decoders.

However, it is important to note that the specificity of NiCLIP’s predictions is bounded by the ontology used for decoding. Although the framework could in principle be extended to predict finer-grained affective states, such as specific emotions, the current Cognitive Atlas vocabulary and task mappings do not provide sufficient coverage for validated emotion-specific decoding. Future work could incorporate specialized affective ontologies to evaluate NiCLIP at this finer level of granularity.

### 3.3. Recommendations, potential use cases, and applications

The results presented in this work highlight the importance of using the full text of the publication for CLIP models, as additional text offers more distinctive context for the associated brain maps. More importantly, we recommend using domain-specific LLMs through fine-tuning (i.e., BrainGPT-7B-v0.2), since they have been shown to produce better text-to-image associations than more general models. Additionally, prediction must rely on accurate and enhanced ontologies, such as the reduced and curated Cognitive Atlas ontology. The current trained model should not be used to decode images with high noise, such as subject-level activation maps, as our decoding model performs poorly on this type of data.

Additionally, we observed that larger ROIs tend to exhibit a decrease in task selectivity, which aligns with previous findings on the relationship between selectivity and the size of brain regions. However, as noted for smaller ROIs (**Fig. 3**), some regions are indeed highly selective, but others remain low in selectivity regardless of their size relative to the whole brain. Caution is advised when interpreting the probabilities of NiCLIP predictions from this kind of predictive model. With the current training data, a high-probability target activation region for a task suggests that the semantic context of the task name and definition closely matches the semantic content of the brain region when mapped into the shared latent space, relative to other task names from the provided ontology. However, this does not mean that directly performing a forward inference experiment—executing the predicted task in an MRI—will necessarily result in selective activation of that specific region alone.

NiCLIP performs functional decoding on whole-brain statistical maps, which provide dense activation patterns across the brain, as well as sparse brain images, such as regions of interest. These capabilities suggest potential applications of reverse inference methods for generating candidate functional annotations of brain regions. However, some predictions warrant interpretive caution. For example, NiCLIP’s prediction of language-related associations for the striatum illustrates how the framework can produce non-obvious hypotheses about regional function. The Language task (“Story vs. Math”) was primarily associated with “visual object recognition” (27.4%) and the Social cognition task with “auditory sentence recognition” (58.0%), neither of which captures the core process being contrasted. Such cases likely reflect the co-occurrence of these paradigms with visual and language processing in the training literature and the task-to-concept structure of the Cognitive Atlas, and indicate that NiCLIP predictions should be treated as candidate hypotheses rather than definitive characterizations. Similarly, the hippocampal ROI was primarily associated with the “working memory fMRI task paradigm” rather than episodic memory, likely reflecting frequent hippocampal reports in n-back studies and the breadth of the composite ROI. Here the domain-level prediction (“Learning and Memory”, 47.2%) is more consistent with the region’s canonical role, underscoring that task-level predictions should be read alongside domain-level outputs and prior literature. The goal of this work is therefore to introduce an accessible decoding framework that researchers can use to generate, evaluate, and refine such hypotheses in future studies. Other applications include leveraging NiCLIP to describe functional domains of regions of interest that may be relevant to a specific population. Such predictions can also be used to contrast the functional associations linked to a region between healthy controls and clinical groups. In summary, the application of reverse inference is not limited to brain activation maps alone. In principle, one could explore the functional domain of any region in the entire brain.

In this work, NiCLIP was trained and validated to predict fMRI tasks, concepts, and domains from brain activation images using a cognitive ontology. However, the proposed framework can be applied to any image and text pairs, as long as the appropriate training data is available. For example, using data properly annotated with unstructured text by trained neuroradiologists, NiCLIP could be used with structural MRI, computed tomography scans, or other medical imaging modalities to identify names of pathologies, structural abnormalities, and other conditions. Note that the framework for the contrastive model presented here only requires embeddings from different sources as input; in principle, a contrastive model could be trained to also predict text related to resting state connectivity and other neuroimaging derivatives data. The network architecture and parameters may need to be adjusted for those applications.

### 3.4. Limitations and challenges

We note several limitations in this current work. The most significant weakness of our study pertains to the small size of the training sample. Typically, CLIP models are trained with hundreds of millions of image-text pairs (Radford et al., 2021). In contrast, our CLIP model was trained with just over 20,000 pairs, which currently is the largest sample of neuroscience papers with complete text information and reported activation coordinates. It is worth noting that (1) brain statistic images are much more constrained than naturalistic images, and (2) we employed a representation of the images based on immense quantities of training data, defined by the DiFuMO atlas. These two factors may mitigate the smaller volume of training data relative to other CLIP applications, which require multiple variations of an image’s content and advanced convolutional neural networks to extract image features. In addition, since NiCLIP was trained primarily on coordinates from positively activated maps, predictions based on negative activation values should be interpreted cautiously. Future training data that include negative activations would be needed to validate decoding of deactivation patterns.

We note a contrast in performance: image-based multi-class classification methods can identify HCP task conditions from individual fMRI images with greater than 90% accuracy (e.g., via linear SVM or deep learning on full statistical maps) (Menuet et al., 2022), whereas NiCLIP achieves a Recall@4 of only 38.19% at the task level for the same subject-level maps. This gap indicates that coordinate-based meta-analytic decoding currently lags behind image-based approaches in individual-level inference, largely because the present model is trained on parcellated CBMA representations rather than full-resolution statistical maps. The primary practical application of NiCLIP in its current form is therefore group-level decoding, where performance is considerably stronger and where the coordinate-based representation more faithfully captures the underlying neural signal. As noted below, extending the training set to include unthresholded statistical maps and subject-level labeled images is a promising route to improving individual-level performance.

The sample size is expected to increase in the future as more articles are published. However, the scientific literature is still growing at a pace that prevents us from achieving a sample size in the millions anytime soon (Bornmann and Mutz, 2015). In principle, we can integrate NeuroVault images into the training set (Gorgolewski et al., 2015). Combining modeled activation maps from peak coordinates with statistical maps has been successfully attempted before (Meudec et al., 2022), yielding improved performance for decoding tasks. We could also incorporate subject-level labeled images from major data consortia, such as ABIDE (Di Martino et al., 2017, 2014), ABCD (Casey et al., 2018), HCP (Barch et al., 2013; Smith et al., 2013; Uğurbil et al., 2013; Van Essen et al., 2013, 2012), ADNI (Jack Jr. et al., 2024), and ENIGMA (Thompson et al., 2014), among others. One could apply different data augmentation techniques and include various versions of the same activation brain map with added noise, or with different thresholds or smoothing parameters. Collectively, these images could expand our sample size to hundreds of thousands. However, we must consider that our CLIP model seems to perform better when the text associated with the images is large. Still, it remains uncertain whether increasing the sample size, even with less detailed text information, can overcome such limitations. More importantly, including subject-level maps and other images from online repositories, such as NeuroVault, could potentially address another issue, namely, the lack of reliability of NiCLIP in predicting individual subject activation maps. Further analyses are necessary to test these hypotheses.

A related limitation concerns how long-form article text is represented. Because full articles often exceed the context window of the LLM encoder, we summarized each article using aggregated chunk-level embeddings rather than a full sequence of word-by-word or paragraph-by-paragraph representations. This mean-pooled document representation is well suited for capturing broad article-level semantic content, such as the cognitive tasks, brain systems, and experimental paradigms discussed in a paper, but it cannot fully preserve the temporal structure, rhetorical flow, or fine-grained dependencies within the text. Future extensions of NiCLIP could use richer document representations, such as attention-weighted pooling, hierarchical encoders, multi-vector retrieval embeddings, or separate embeddings for methods, results, and discussion sections. Such approaches may preserve more nuanced semantic information while retaining the scalability needed for large neuroimaging corpora.

Another limitation of the current implementation is the mismatch between the text used during CLIP training and the text used during ontology-based decoding. The contrastive model was trained using long article-level text, whereas decoding relies on relatively short Cognitive Atlas task names and definitions. This difference may introduce a distributional shift between the training text embeddings and the ontology-entry embeddings used at inference, limiting task-level decoding accuracy. Recent work on NeuroConText has highlighted a related issue and proposed LLM-based text augmentation to improve generalization to short-text inputs (Ghayem et al., 2026). Future versions of NiCLIP could similarly enrich Cognitive Atlas entries by expanding task and concept definitions into longer, more publication-like descriptions or by retrieving representative passages from the neuroimaging literature. Such ontology enhancement may reduce the mismatch between training and inference text and improve fine-grained decoding performance. This text mismatch may be especially consequential for subject-level activation maps, where lower signal-to-noise ratio and individual variability further reduce the reliability of the image embedding and make it harder to match activation patterns to short ontology descriptions. Thus, the weaker subject-level performance likely reflects both image-level variability and the train–inference text mismatch, rather than image noise alone. Future work should more directly evaluate the effect of signal-to-noise ratio on decoding performance by constructing group-average maps from increasing numbers of subjects and testing whether NiCLIP predictions improve as maps become more stable. Such an analysis would help distinguish the contribution of image-level noise from the train–inference text mismatch introduced by comparing long article-level embeddings with short ontology descriptions.

In addition, the formulation of the reverse inference problem follows Bayes’ theorem, where the posterior probability of a task name given an activation is directly proportional to its prior probability. Here, we have chosen to set the prior distribution on tasks based on their prevalence in the literature. However, this assumption may be suboptimal for rare tasks, and a uniform prior could be used as an alternative. Additional work is needed to assess the actual influence and provide recommendations for selecting prior probabilities.

Finally, although NiCLIP was evaluated on HCP group-level maps, subject-level maps, and ROI maps, broader cross-dataset validation remains an important next step. The framework should in principle be applicable to independent task-fMRI datasets when their contrasts can be mapped onto the decoding ontology, but performance is expected to depend on the similarity between the input maps, the training literature, and the decoding vocabulary. Datasets that include tasks that differ from the HCP battery may prove useful as future benchmarks. For example, preliminary investigation of NiCLIP’s generalizability to the Individual Brain Charting (IBC) dataset (Pinho et al., 2018), which includes tasks similar to the HCP, as well as additional spatial, social, and emotional tasks, demonstrated consistent performance in terms of task-level decoding accuracy (**Fig. S10**). In contrast, the Natural Scenes Dataset (NSD; (Allen et al., 2022)) may primarily test visual-perception decoding and may underperform for scene-specific content because naturalistic scene viewing is not well represented in the current coordinate-based training corpus.

## 4. Conclusions

Taken together, this study presents NiCLIP, the first LLM-powered CLIP model for formal reverse inference of brain activation maps. We confirmed CLIP’s unique ability to match article text to brain images. The results indicate that a fine-tuned LLM provides the most effective text-to-brain association, with generally stronger performance when using the full body of the articles. We validated NiCLIP as a precise decoding tool for group-level dense activation maps; however, our decoding model underperformed on individual subject images. Additionally, NiCLIP demonstrated notable performance with other image types, such as ROI maps. Lastly, we highlighted the importance of improving cognitive ontologies for reverse inference tasks. NiCLIP extends earlier methods for describing activation maps and may support future hypothesis generation by providing structured, ontology-guided candidate interpretations of brain activation patterns (Poldrack, 2006).

## 5. Methods

### 5.1. Datasets

#### 5.1.1. PubMed neuroimaging papers

We conducted an extensive search for neuroimaging articles in PubMed Central using Pubget (https://neuroquery.github.io/pubget/pubget.html), an open-access Python tool for collecting data for biomedical text mining. We performed a query and retrieved articles that included keywords such as “fMRI” or “functional MRI” in the abstracts (e.g., “(fMRI[Abstract]) OR (functional MRI[Abstract])”). Pubget extracted text, metadata, and stereotactic coordinates from the articles in a standardized format. The data was then converted into a NiMARE database object to facilitate downstream tasks such as term-based and topic-based meta-analysis for the baseline models and to generate images from the extracted activation coordinates. As of February 2025, the PubMed search for neuroimaging articles yielded more than 30,000 papers. A total of 23,865 articles included at least one activation coordinate, a complete abstract, and full-text information.

The text embeddings *Mε*ℜ^23,865×4,096^ were derived using pre-trained LLMs. We compared four different LLM models: BrainGPT models (i.e., BrainGPT-7B-v0.1 and BrainGPT-7B-v0.2) (Luo et al., 2024), along with their corresponding foundational pre-trained LLM (i.e., Llama-2-7b-chat-hf and Mistral-7B-v0.1, respectively) (Jiang et al., 2023; Touvron et al., 2023). Due to the token size limitation of these LLMs, embeddings for long text were generated by averaging the embeddings from smaller chunks of the text. Specifically, each article was divided into chunks that fit within the context window of the corresponding LLM encoder. A separate embedding was computed for each chunk, and the final article-level embedding was obtained by averaging the chunk embeddings. This procedure produces a document-level semantic summary of the article rather than a full sequence representation of the text. Article-specific brain images were generated from brain activation coordinates using a Multilevel Kernel Density Analysis (MKDA) (Wager et al., 2007). MKDA is a kernel-based method that convolves each activation coordinate with a binary sphere of a set radius around the peak voxel. The coordinate-specific binary maps (i.e., one map corresponding to each coordinate from the set) were combined into a single modeled activation map by taking the maximum value for each voxel. In this work, we used a sphere with a radius of 10mm. Finally, brain image embeddings *Aε*ℜ^23,865×512^ were extracted from the modeled activation map using a continuous brain parcellation defined by the DiFuMo atlas with 512 regions (Dadi et al., 2020). Text and image embeddings were L2 normalized. We used DiFuMo512 to match the NeuroConText framework and to balance spatial resolution with model complexity. DiFuMo1024 would double the dimensionality of the image embedding, increasing computational cost and potentially requiring additional architectural tuning.

#### 5.1.2. Cognitive Ontology

Cognitive Atlas (Poldrack et al., 2011) (https://www.cognitiveatlas.org/) is an online repository of cumulative knowledge from experienced researchers in psychology, cognitive science, and neuroscience. The repository currently offers two knowledge bases: 912 cognitive concepts and 851 tasks, complete with definitions and properties. The cognitive concepts establish relationships with other concepts and tasks, aiming to create a map between mental processes and brain functions. The task and concept metadata (i.e., names and definitions) were downloaded using the Cognitive Atlas API, while the task, concept, and domain relationships were obtained through NiMARE’s ‘download_cognitive_atlas’ function. Here, a “domain” refers to a high-level concept, as defined by the concept categories in Cognitive Atlas (https://www.cognitiveatlas.org/concepts/categories/all).

Despite the extensive collection of tasks and concepts, the Cognitive Atlas mapping remains limited. Although the full Cognitive Atlas contains 851 tasks and 912 concepts, many entries have incomplete definitions, missing task-to-concept mappings, or sparse concept-to-domain annotations. Additionally, as a community-based ontology, these mappings reflect the opinions of individual researchers and may not always be widely agreed upon or empirically validated. These limitations can directly affect ontology-guided decoding because task-level predictions are propagated to concepts and domains through these mappings. To reduce this source of noise, we used the reduced Cognitive Atlas ontology derived from Menuet et al. (Menuet et al., 2022), which provides a curated subset of commonly used fMRI tasks and more complete task-to-concept mappings. Task-level predictions were evaluated against the corresponding Cognitive Atlas task labels, and concept- and domain-level predictions were derived from the ontology’s task-to-concept and concept-to-domain relationships. Thus, the reduced ontology is both smaller in vocabulary size and more complete in its task-to-concept annotations.

#### 5.1.3. Task fMRI datasets

The functional decoding model was assessed using the task-fMRI group-average maps from the HCP S1200 data release (Barch et al., 2013; Smith et al., 2013; Uğurbil et al., 2013; Van Essen et al., 2013, 2012). Specifically, we used the results in volume space reported in collection 457 in NeuroVault (https://neurovault.org/collections/457/). The HCP tasks target seven major domains that sample a diverse set of neural systems of high interest in the neuroscience field. These domains include (1) emotion processing, (2) category specific representations, (3) language processing (semantic and phonological processing), (4) visual, motion, somatosensory, and motor systems, (5) relational processing, (6) social cognition (theory of mind), (7) working memory/cognitive control systems. Collection 457 contains multiple contrast maps across these seven HCP domains. For task-level decoding, we selected one representative contrast per domain that best captured the core cognitive process of the task: Faces vs. Shapes for Emotion, Reward vs. Baseline for Gambling, Story vs. Math for Language, the average of all movement contrasts for Motor, Relational vs. Match for Relational processing, TOM vs. Random for Social cognition, and 2-Back vs. 0-Back for Working Memory. These seven representative contrasts were used as the task-level evaluation maps in the group-level HCP decoding analysis. Additional details regarding these tasks and their available contrasts have been previously published (Barch et al., 2013). Next, the functional decoding model was evaluated using subject-level statistical images, as these data tend to have more variability and noise compared to group-level images. We used data from subject-level activation maps from 787 individuals, which contain the same tasks and contrasts as those in the HCP group-level analysis.

We note that the PubMed training corpus may include articles analyzing HCP data. However, several factors mitigate potential leakage. First, the CLIP model was trained on MKDA-modeled coordinate maps derived from reported activation coordinates, which are structurally distinct from the group-level HCP t-statistic maps used for evaluation. Second, even if an article discusses an HCP contrast, the text embedding captures the full article content rather than a one-to-one mapping to a specific statistical map. Third, cross-validation folds were constructed by randomly sampling articles without topic stratification, reducing the risk that a specific task domain was systematically isolated in one fold. Thus, although text-level overlap with HCP-related publications cannot be fully excluded, direct image-level leakage from the HCP evaluation maps is unlikely.

#### 5.1.4. Meta-analytic regions of interest

Next, the functional decoding model was evaluated on specific regions of interest (ROI). We used six highly popular anatomical brain regions, including the amygdala (Bzdok et al., 2013a), hippocampus (Plachti et al., 2019), insula (Chang et al., 2013), striatum (Liu et al., 2020), right temporoparietal junction (rTPJ) (Bzdok et al., 2013b), and ventromedial prefrontal cortex (vmPFC) (Chase et al., 2020). Here, we defined a single ROI for each brain region by combining all the seed regions. **Fig. S11** presents the seed regions per ROI as defined by their corresponding meta-analytic-based parcellation analysis.

### 5.2. CLIP model for article and brain image association

The representation-learning component of NiCLIP intentionally follows the NeuroConText framework closely (Ghayem et al., 2026). Specifically, we use the same general CLIP-style strategy of aligning text embeddings derived from articles with brain embeddings derived from coordinate-based modeled activation maps. In this way, the novel methodological contribution of NiCLIP lies primarily in the downstream ontology-guided decoding framework. The CLIP model architecture (Fig. 4A) adheres to the identical settings employed in the NeuroConText framework (Ghayem et al., 2026). CLIP consists of both a text and an image encoder. The text encoder consists of a projection block and two residual blocks, while the image encoder comprises three residual blocks. Each block contains a fully connected layer, GELU activation, dropout, and layer normalization. We use ‘block’ rather than ‘head’ to distinguish these architectural components from the attention heads in transformer models. The projection block takes the high-dimensional text embedding and projects it into a shared dimensional space to align with the image embedding dimension, which has already been reduced through brain parcellation. The projection block consists of a fully connected projection layer, followed by a GELU activation function, an identity fully connected layer, a dropout rate of 0.5 for regularization, and a normalization layer. Each residual block includes an identity fully connected layer and a GELU activation function, followed by a dropout rate of 0.5 and a normalization layer.

**Figure 4.**
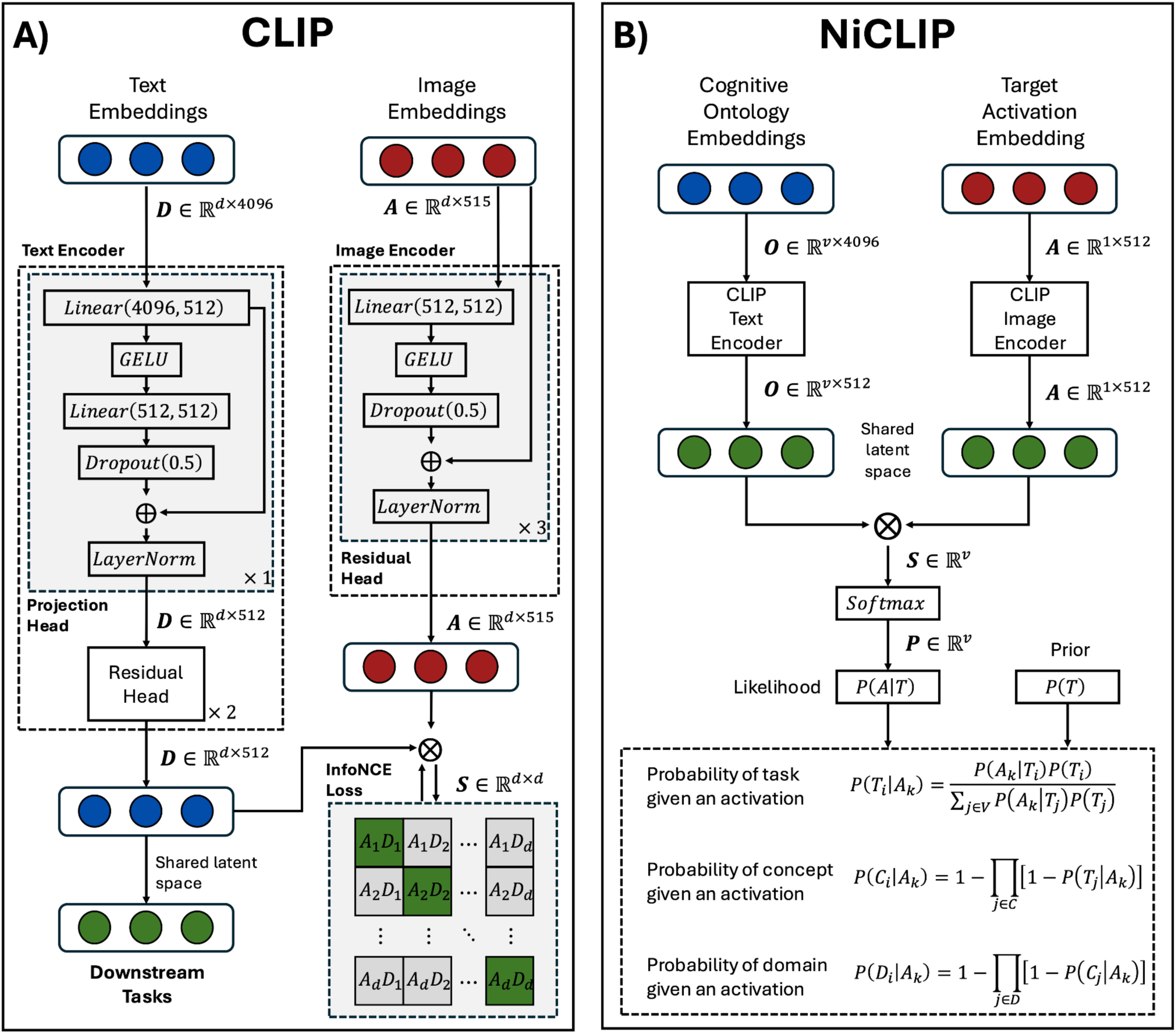
CLIP and NiCLIP model architecture. **(A)** The architecture of the CLIP model includes a text encoder and an image encoder that transform input embeddings into a shared latent space. The text encoder consists of a projection block and two residual blocks, while the image encoder has three residual blocks. The projection block is defined by a linear projection layer, followed by a GELU activation function, a linear layer, a dropout layer, and culminating in a normalization layer. The residual block is made up of a linear identity layer, followed by a GELU activation and a dropout layer, concluding with a normalization layer. The output from the shared latent space is utilized for downstream tasks (e.g., functional decoding), and InfoNCE loss is applied in the latent space for self-supervised learning during training. **(B)** NiCLIP takes advantage of the task name embedding and the extracted features from a target activation map. These embeddings are encoded with the pre-trained CLIP text and image encoders. Cosine similarity is assessed in the shared latent space, and a softmax function converts them to a likelihood P(A|T). Using the prior probability P(T), we compute the posterior probability of a task given an activation P(T|A). Furthermore, the noisy-OR model is employed to determine the probability of a concept P(C|A) and domain P(D|A) given an activation.

The model’s hyperparameters for training were selected from the NeuroConText paper (Ghayem et al., 2026), with the following configurations: batch size of 128, learning rate of 5e-4, weight decay of 0.1, and a total of 50 epochs. The contrastive loss function InfoNCE was employed for self-supervised learning during training, following the standard implementation of the CLIP model (Oord et al., 2019). Additionally, we incorporated an early stopping rule to prevent overfitting and avoid unnecessary computation during epochs without performance improvement. We monitored the validation loss across epochs while maintaining a patience window of 10 epochs to accommodate potential performance fluctuations. If the validation loss did not improve within the patience window, training was terminated, and the model weights corresponding to the lowest validation loss were saved. Finally, the model was trained and evaluated using distinct data loaders for training, validation, and testing, implementing a 23-fold cross-validation on the test set and a 22-fold nested cross-validation on the evaluation set. The fold size for the testing and evaluation was 1,000 samples, resulting in approximately 21,865 articles available for training.

### 5.3. NiCLIP: neuroimaging CLIP model for predicting tasks, concepts, and domains from brain images

The NiCLIP architecture (**Fig. 4B**) leverages the learned shared latent space from the CLIP model to make predictions on unseen data. Given a target activation map, the goal of the NiCLIP decoder is to predict the most likely task, concept, and domain associated with the activation pattern. NiCLIP relies on a cognitive ontology to define the fMRI tasks and their definitions, as well as their associations with concepts and domains. Using Bayes’ theorem, we can express the probability that a brain activation pattern (*A_k_*) was produced by a certain fMRI task *T_i_* as:

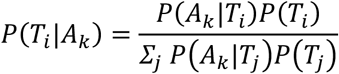

where *P*(*T_i_*|*A_k_*) is the posterior probability, *P*(*A_k_*|*T_i_*) is the likelihood of task *T_i_* having an activation pattern *A_k_*, and *P*(*T_i_*) is the prior probability of task *T_i_* regardless of the activation pattern.

First, we use the pre-trained CLIP model to estimate the likelihood *P*(*A_k_*|*T_i_*). To accomplish that, we use a vocabulary composed of all Cognitive Atlas tasks and their definitions. Initially, the vocabulary was encoded by the same LLM used to train CLIP, and then the embedding for each task was determined as a linear combination of the task name embedding and its definition:

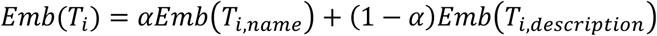

where the parameter *α* represented the weight of the name embedding relative to its definition. Here, we considered *α* = 0.5.

Next, the vocabulary embeddings were encoded by the text encoding layers from CLIP, projecting the vocabulary into the shared latent space with the images. For the target image, we extracted 512 features using the Difumo atlas (i.e., the same dimension used to train CLIP). We encoded the features with the image encoding layers, projecting the images into the same shared latent space of the vocabulary. Finally, we calculated the cosine similarity between the vocabulary embeddings and the input image embeddings and transformed the similarity into probability, resulting in the likelihood *P*(*A_k_*|*T_i_*). We note that *P*(*A_k_*|*T_i_*), as defined by the softmax of cosine similarities in the CLIP latent space, is a heuristic likelihood that quantifies the relative compatibility between activation patterns and task descriptions. Although this normalized score is non-negative and sums to one across candidate tasks, it is not calibrated as a proper statistical likelihood. Similarly, the posterior *P*(*T_i_*|*A_k_*) should be interpreted as a relative ranking score rather than a rigorously calibrated probability.

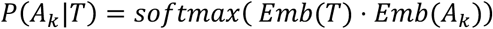

The prior probability *P*(*T_i_*) was determined by the task representation in the neuroscience corpora used for training the CLIP model. The task representation was defined by the cosine similarity of all document task pairs DT. To obtain a measure of the global representation, we calculated the mean across all publications, resulting in a similarity score for each task with the corpora. Finally, the prior probability resulted in

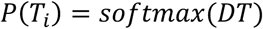

Now, since we have the posterior probabilities of the task given an activation P(T|A), we can use noisy-OR model to define the probability of a concept (*C_j_*) given an activation *A*_*k*_ as:

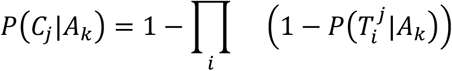

where 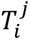 represents tasks that measure the concept *C_j_*. The noisy-OR model used to compute *P*-*C_j_*|*A_k_* assumes conditional independence among tasks that measure the same concept. This assumption is a simplification, because tasks sharing a concept may have correlated activation patterns and overlapping semantic content. The resulting concept and domain probabilities should therefore be interpreted as approximate scores that aggregate evidence across related task predictions, rather than as calibrated posterior probabilities. Similarly, we can obtain the probability of the domain *D_j_* given an activation *A_k_* as:

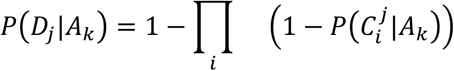

where 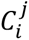 represents concepts categorized under the cognitive process *D_j_*.

### 5.4. Evaluation metrics

We assessed the CLIP model’s ability to match brain images with their corresponding text using Recall@k and Mix&Match (Menuet et al., 2022; Ghayem et al., 2026; Mitchell et al., 2008), which are commonly employed in contrastive learning models. Given an activation map, Recall@K quantifies the likelihood of finding its true corresponding text among the top-k ranked by similarity within the entire sample (Ghayem et al., 2026). To determine the top-k maps, we first calculate the similarity matrix *Sε*ℜ^n×n^ between the image and text embeddings in the shared latent space. The identity matrix represents the true positive, as the diagonal element indicates the similarity between an image and its corresponding text. As specified in the InfoNCE loss, the model is trained so that the diagonal element of the similarity matrix approaches 1 while the nondiagonal elements decrease towards zero. The similarity matrix is sorted on a row-by-row basis, and we check whether the diagonal element (i, i) is among the top-k ranked elements. The final scores are computed by averaging across rows:

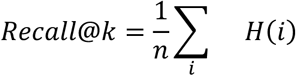

where n represents the total number of samples (or rows), and H denotes the indicator function that yields one if the column j=i belongs to the top k sorted columns (*i ε Top_ki_*) and zero otherwise. Specifically, the best model was determined based on the highest test set Recall@10

Mix&Match measures the likelihood that a given brain map is more similar to its true corresponding text than the other text embeddings in the set; Mitchell et al., 2008). In other words, this metric assesses whether the learned embeddings in the shared latent space representations are discriminative, indicating that an item is more similar to itself than to other items.

We also evaluated the sensitivity of the functional decoders, including baselines and NiCLIP, using Recall@K. In this context, the Recall@K value can be interpreted as the likelihood that a decoder ranks the true label among the top-k labels for a given brain activation map (Menuet et al., 2022). We utilized the image metadata from the HCP, which contains the corresponding Cognitive Atlas task as the ground truth.

## Acknowledgments

Special thanks to the FIU Instructional & Research Computing Center (IRCC, http://ircc.fiu.edu) for providing the HPC and computing resources that contributed to the research results reported in this paper.

## Ethical Statement

The Human Connectome Project provided the ethics and consent needed for the study and dissemination of HCP data. This secondary data analysis was approved by the Institutional Review Board of Florida International University.

## Funding Statement

Funding for this project was provided by NIH R01-MH096906.

## Data Accessibility

The functional neuroimaging data were provided by the Human Connectome Project, WU-Minn Consortium (Principal Investigators: David Van Essen and Kamil Ugurbil; U54-MH091657) funded by the 16 NIH Institutes and Centers that support the NIH Blueprint for Neuroscience Research; and by the McDonnell Center for Systems Neuroscience at Washington University. The results of group-level activation maps in volume space can be downloaded from collection 457 in NeuroVault (https://neurovault.org/collections/457/). The subject-level activation maps can be downloaded from collection 4337 in NeuroVault (https://neurovault.org/collections/4337/).

The meta-analytic parcellation images are publicly available for download at https://anima.fz-juelich.de/.

A simplified representation of the Cognitive Atlas task is available in GitHub https://github.com/Parietal-INRIA/fmri_decoding/blob/master/Data/labels/cogatlas_tasks_concepts_bertrand_mini.csv.

## Code Availability

This project relied on multiple open-source Python packages, including: Jupyter (Kluyver et al., 2016), Matplotlib (Hunter, 2007), Neuromaps (Markello et al., 2022), NiBabel (Brett et al., 2020), Nilearn (Abraham et al., 2014), NiMARE (Salo et al., 2024, 2023), PyMARE (Yarkoni et al., 2024), NumPy (van der Walt et al., 2011), Pandas (McKinney, 2010), Scikit-learn (Pedregosa et al., 2011), SciPy (Virtanen et al., 2020), Seaborn (Waskom, 2021), and SurfPlot (Gale et al., 2021). We also used the HCP software Connectome Workbench (wb_command version 1.5.0, (Marcus et al., 2011)).

All code required to reproduce the analyses and figures in this paper is available on GitHub at https://github.com/NBCLab/brain-decoder. The *braindec* Python package and documentation, including a user guide, installation instructions, and examples are provided at https://brain-decoder.readthedocs.io/en/latest/. In addition, a Google Colab notebook, which includes a NiCLIP tutorial and examples, is available at https://colab.research.google.com/github/jdkent/brain-decoder/blob/main/docs/auto_examples/02_niclip_demo.ipynb. A browser-based web interface for NiCLIP is available at https://huggingface.co/spaces/jdkent/niclip, allowing researchers to upload a brain activation map in NIfTI format and receive cognitive decoding results without installing any software. As a longer-term community resource, we plan to integrate NiCLIP into Neurosynth Compose (compose.neurosynth.org; (Kent et al., 2026)), a publicly accessible platform for neuroimaging meta-analysis. All data and resources that resulted from this paper are openly disseminated and made available on the Open Science Framework (OSF) at https://osf.io/dsj56/, including the links to the GitHub repository and figures.

## Competing Interests

The authors declare no competing interests.

## Author Contributions

ARL, JAP, AdlV, JBP, TEN, and JDK conceived and designed the project. JAP, AdlV, and JDK analyzed data. JAP and JDK contributed scripts and pipelines. JAP, TEN, and ARL wrote the paper; AdlV, JDK, and ARL contributed to the revisions; all authors approved the final version.

## Notes

https://osf.io/dsj56/

### Competing Interest Statement

The authors have declared no competing interest.

### Summary of Updates

This version reflects revisions made during peer review. A standalone browser-based web interface for NiCLIP is now available, and the Discussion was expanded to clarify that group-level decoding is NiCLIP's primary use at present. We also softened comparisons between LLMs and made minor textual corrections. The datasets, model architecture, and core results are unchanged.

