## Supplemental Information for "NiCLIP: Neuroimaging contrastive language-image pretraining model for predicting text from brain activation images" (from another version of this preprint: version 3, 2026-05-22)

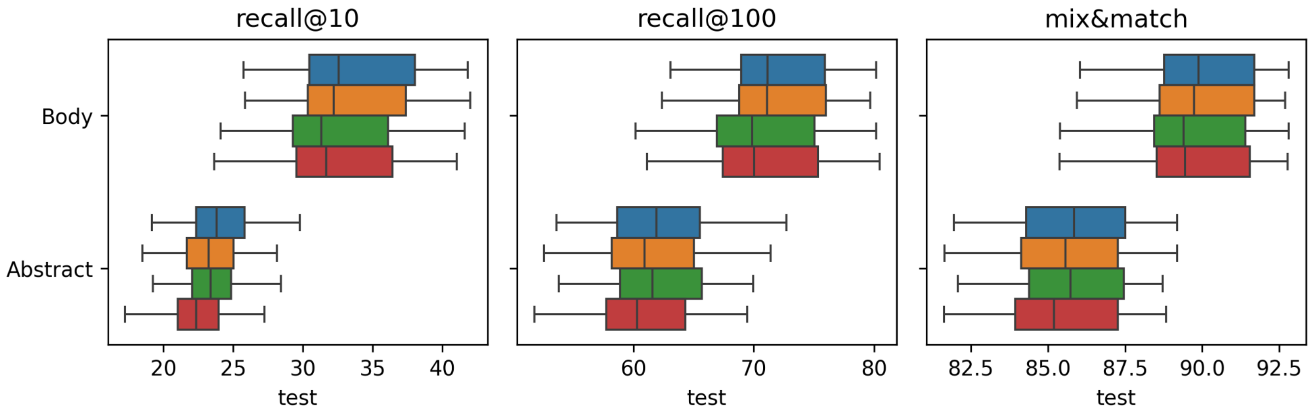


**Figure S1. Text-to-brain association performance comparison.**

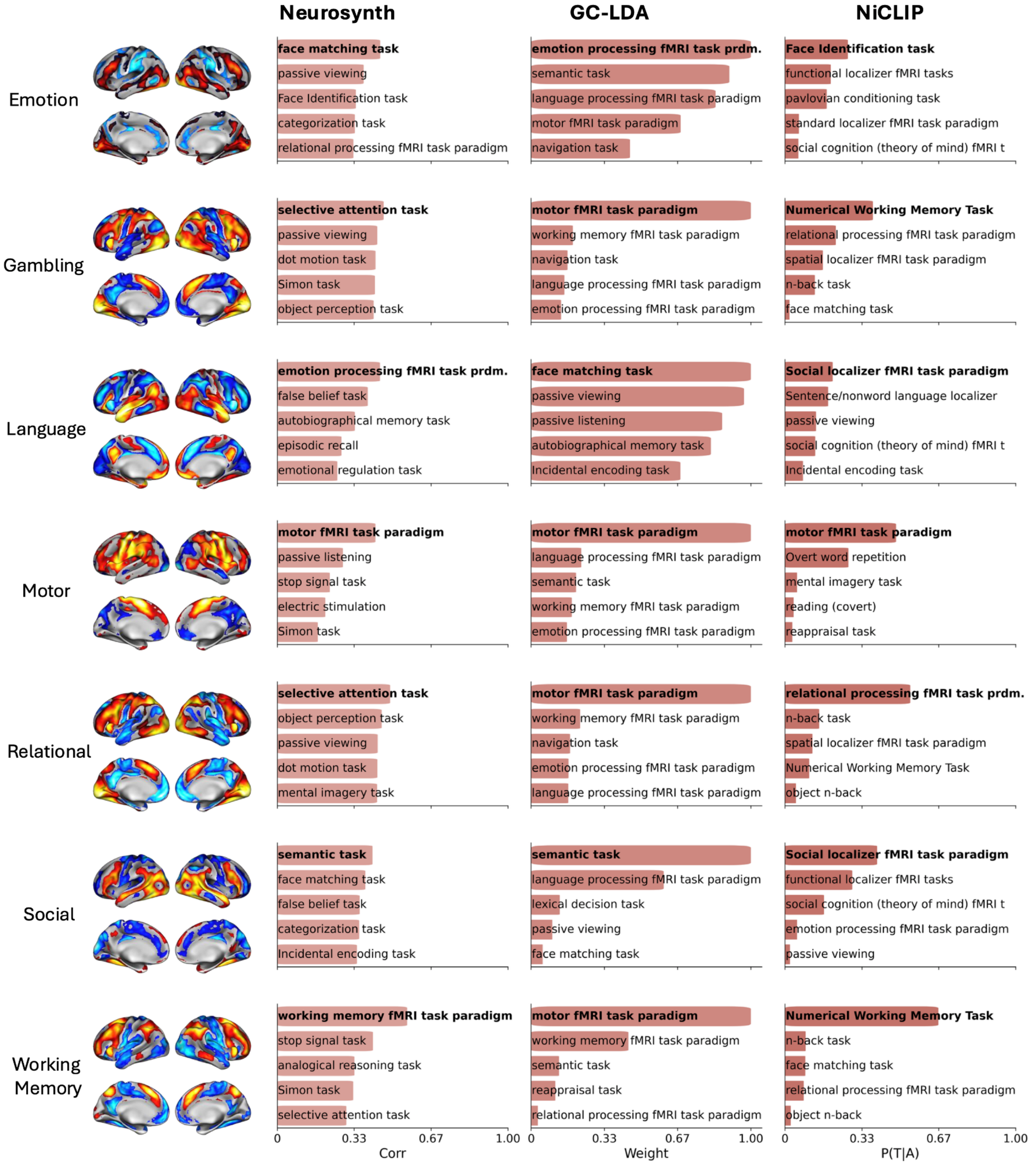


**Figure S2. Comparisons between NiCLIP and baseline models' task predictions with group-level maps.**

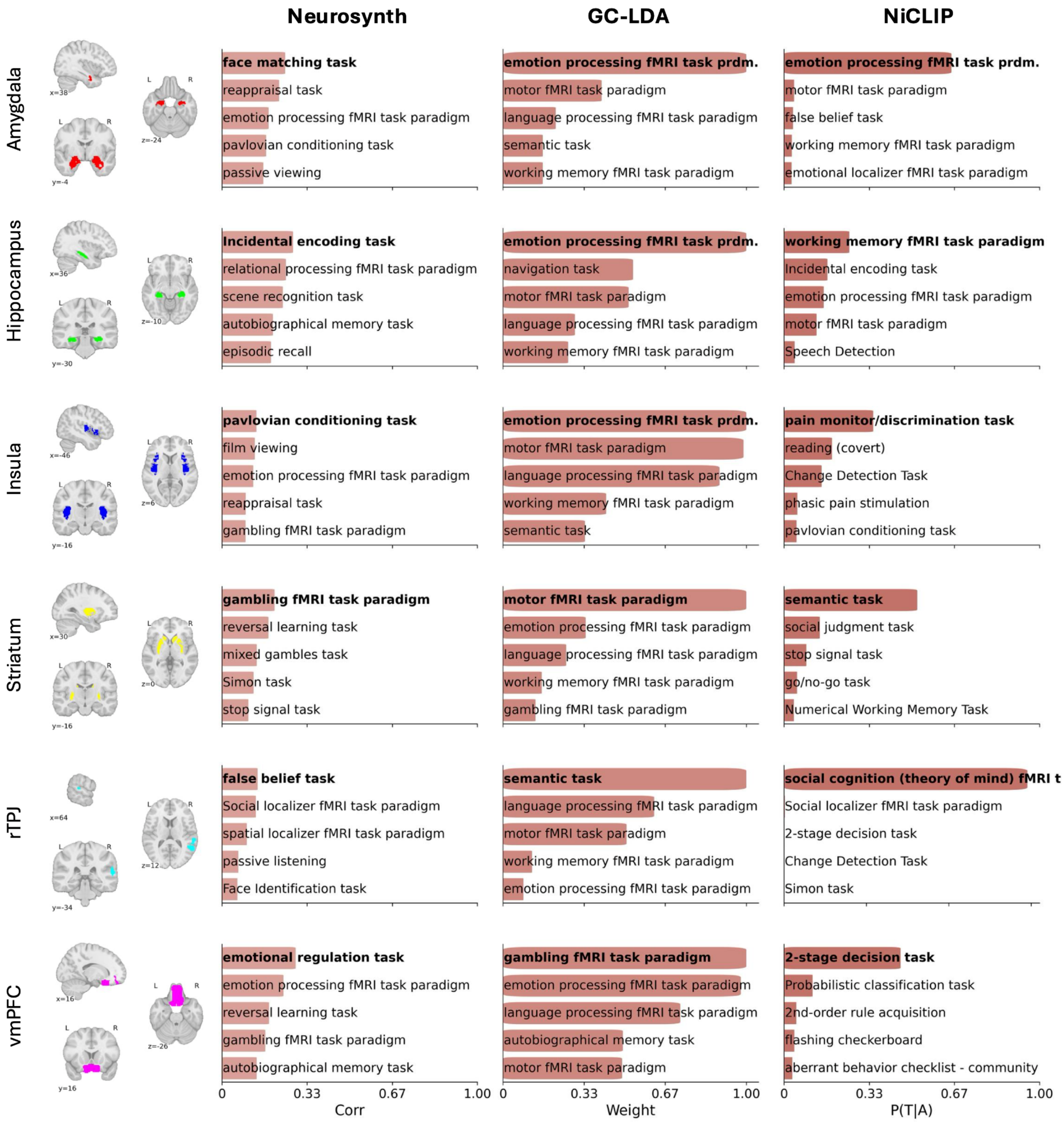


**Figure S3. Comparisons between NiCLIP and baseline models' task predictions with brain ROIs.**

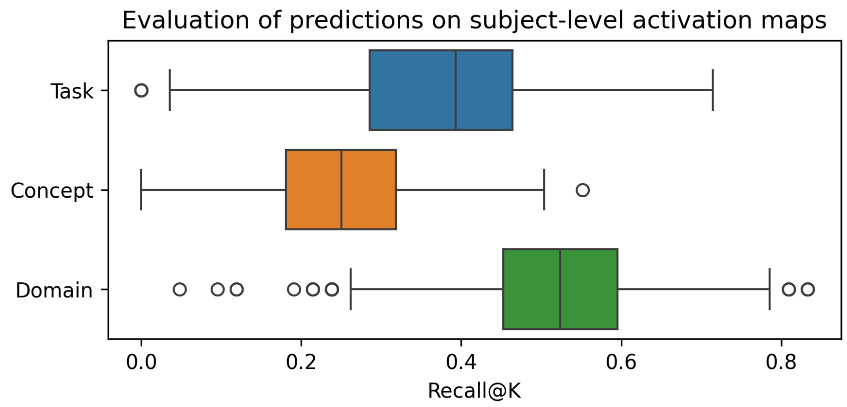


**Figure S4. Distribution of Recall@K score for task, concept, and domain predictions by NiCLIP on subject-level activation maps.**

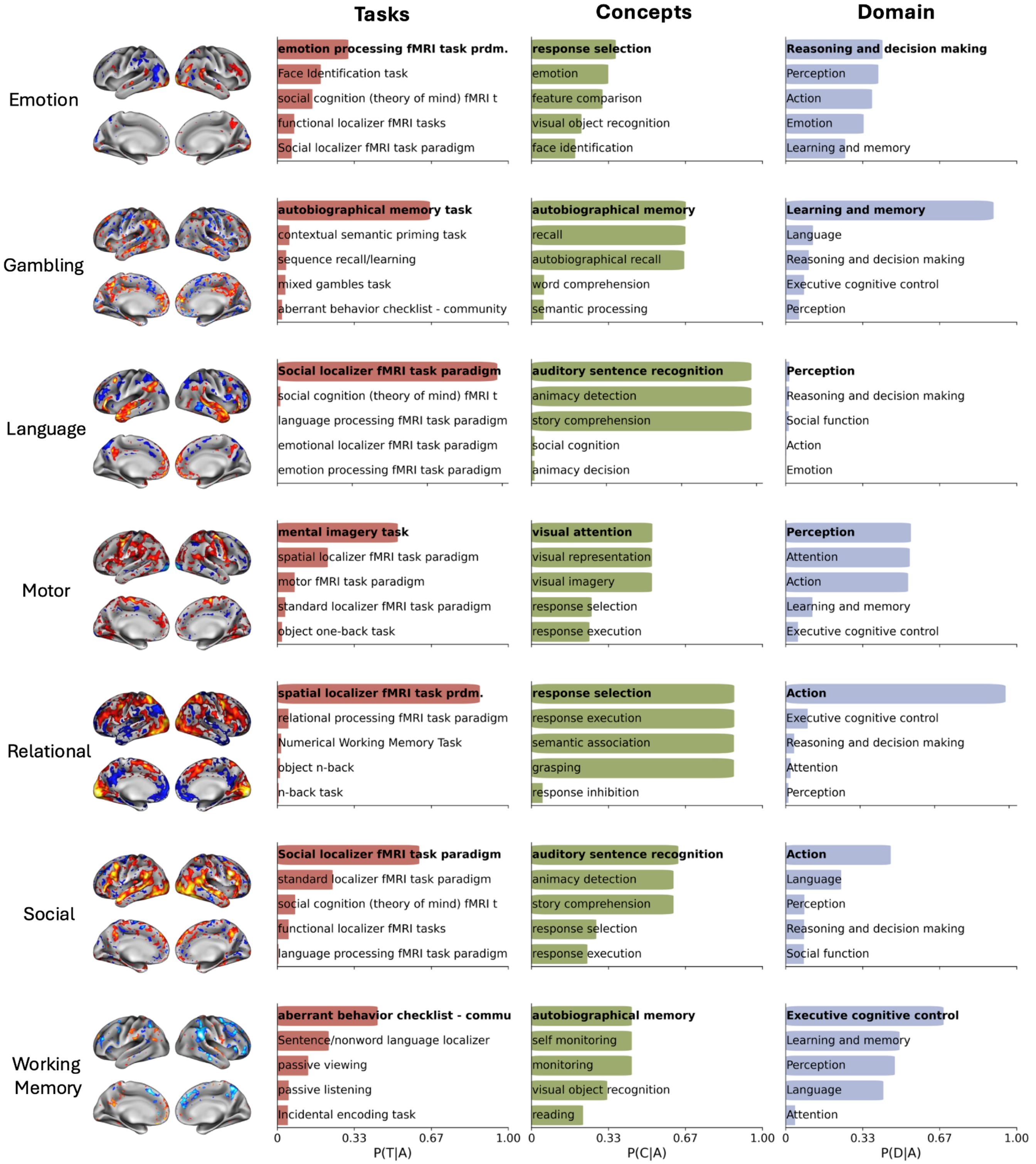


**Figure S5. NiCLIP for predicting tasks, concepts, and domains from brain activation patterns of subject-level maps.**

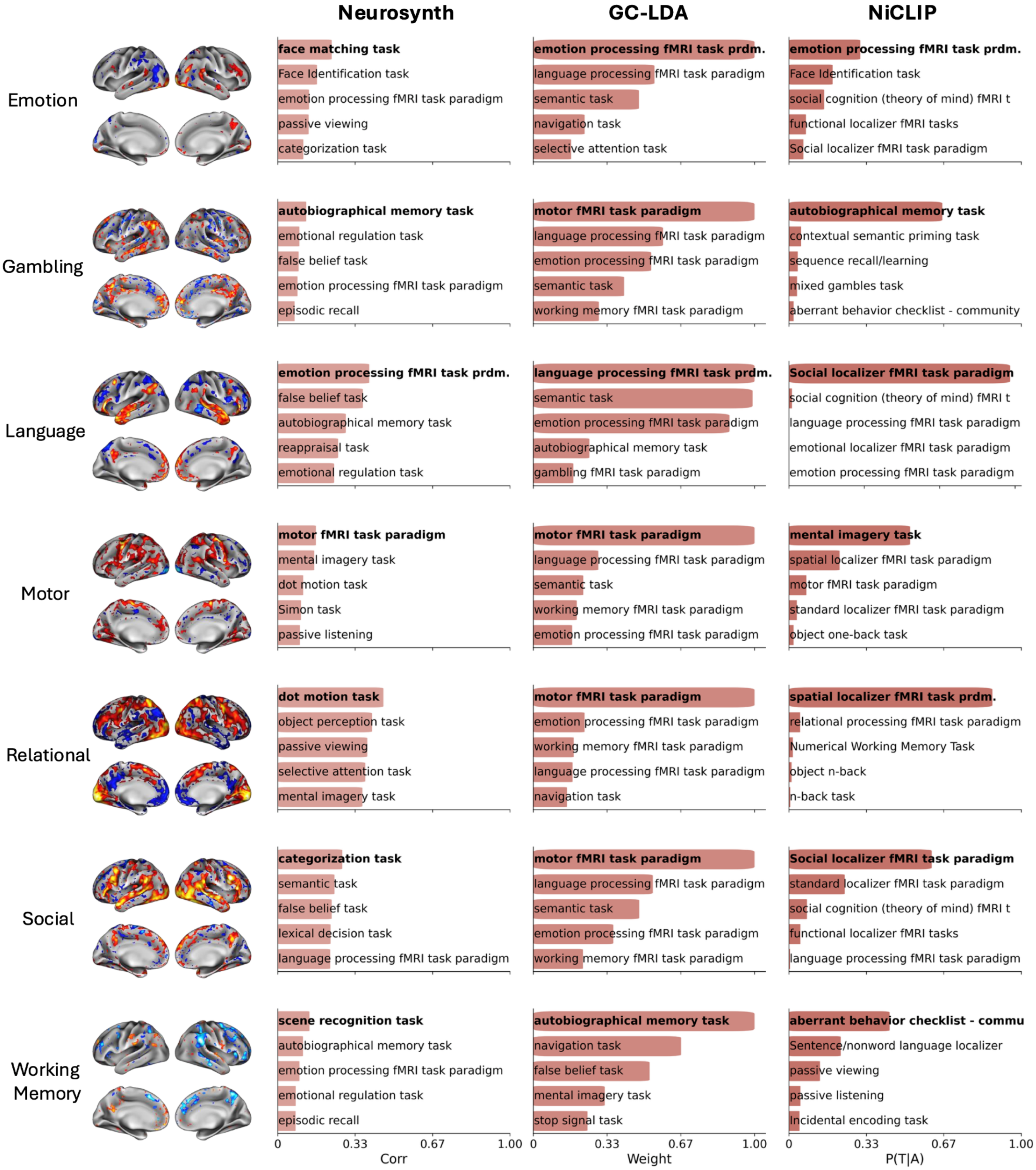


**Figure S6. Comparisons between NiCLIP and baseline models' task predictions with subject-level maps.**

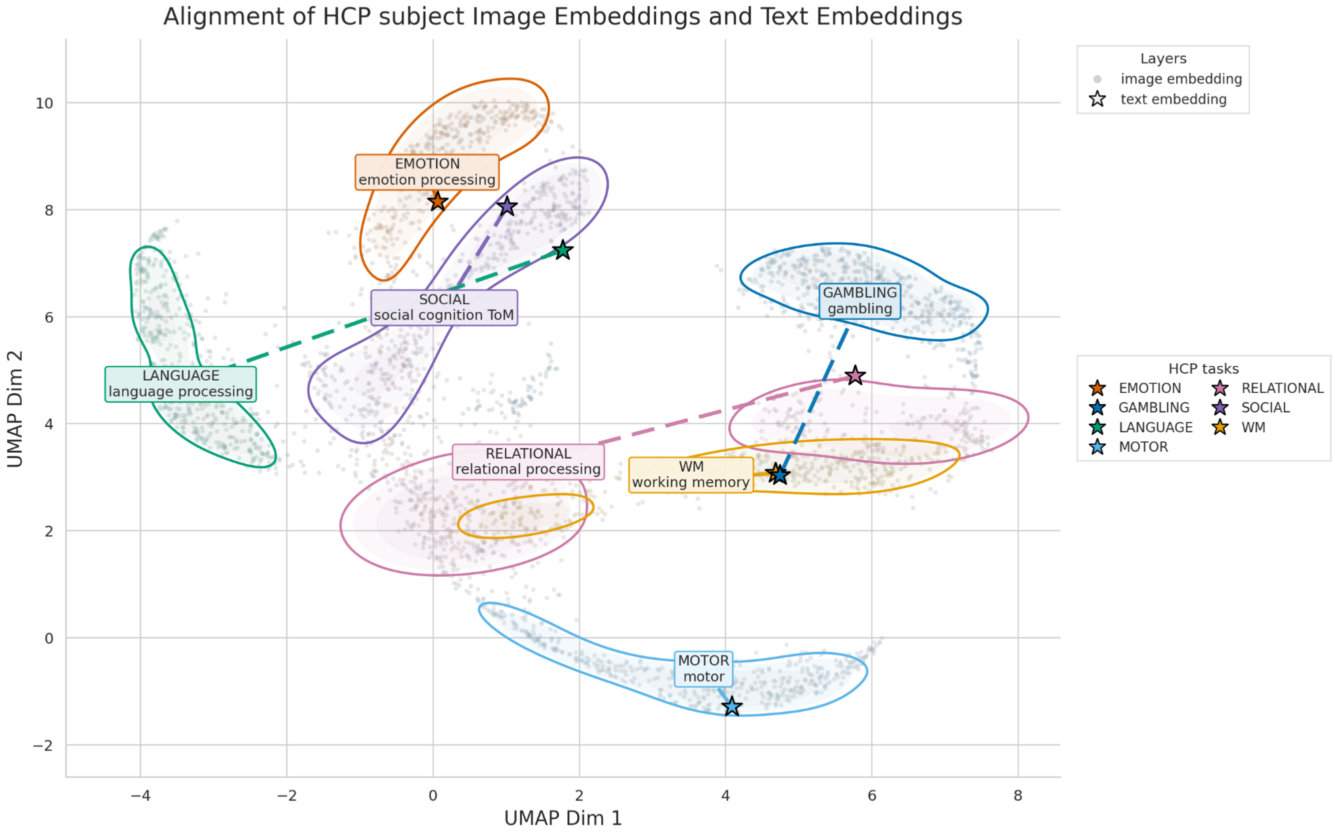


**Figure S7. Alignment of HCP subject image embeddings and text embeddings**. NiCLIP 512-dimensional embeddings were computed for each HCP subject contrast image, then projected to two dimensions using UMAP (n_neighbors=40, min_dist=0.15, cosine metric). The UMAP manifold was fit exclusively on the image embeddings; Cognitive Atlas task text embeddings (generated by BrainGPT-7B-v0.2 from task descriptions) were projected into the same space using the fitted transform, without influencing the manifold geometry. Each grey point represents a single subject contrast image. Colored contours show kernel density estimates of the image distribution for each of the seven HCP task paradigms (i.e., emotion, gambling, language, motor, relations, social, working memory). Colored stars show the corresponding Cognitive Atlas text embedding for each task, with dashed lines connecting each label to the centroid of its image cluster. Close proximity between a star and its cluster centroid indicates strong alignment between the learned brain image representation and the text embedding of the associated task description.

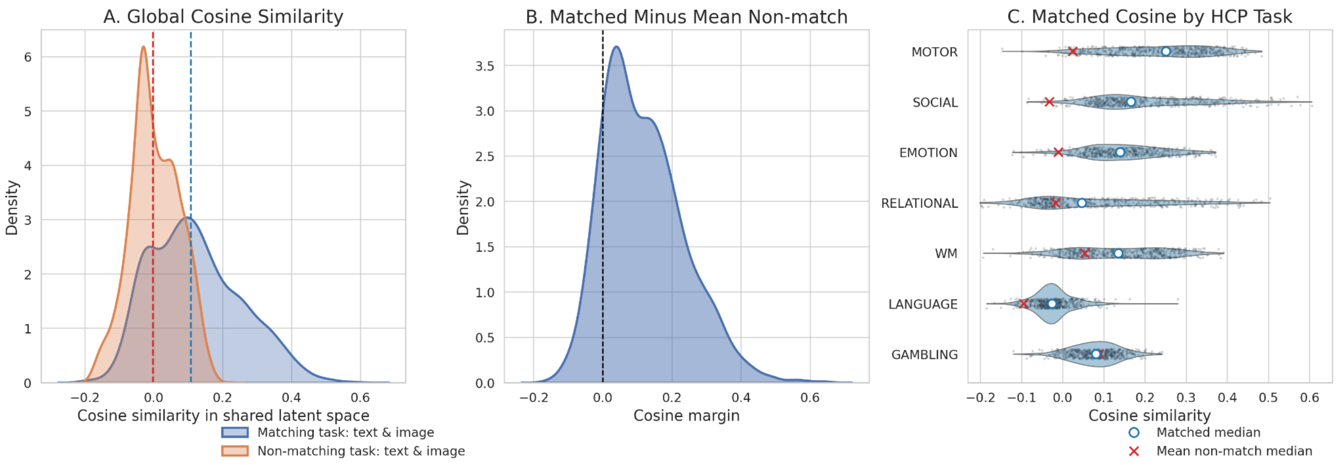


**Figure S8. Cosine similarity between matching image and text embedding**. For each HCP subject contrast image, BrainCLIP embeddings were computed and compared against BrainGPT-7B-v0.2 text embeddings for all Cognitive Atlas task terms via cosine similarity. (A) Global distributions of cosine similarity for matched text–image pairs (image compared against the embedding of its true task label) versus the mean cosine similarity across all non-matching task texts. Dashed lines indicate medians. (B) Distribution of the cosine margin — the difference between matched cosine similarity and the mean non-match cosine similarity — across all subject images. Values greater than zero indicate that a brain image is on average more similar to its correct task text than to other tasks in the vocabulary. (C) Per-task distributions of matched cosine similarity (violin plots with individual subject points), with matched median (open circle) and mean non-match median (red cross) overlaid. Tasks are ordered by mean cosine margin. MOTOR and SOCIAL show the strongest and most consistent text–image alignment, while GAMBLING shows the weakest separation from the non-match baseline.

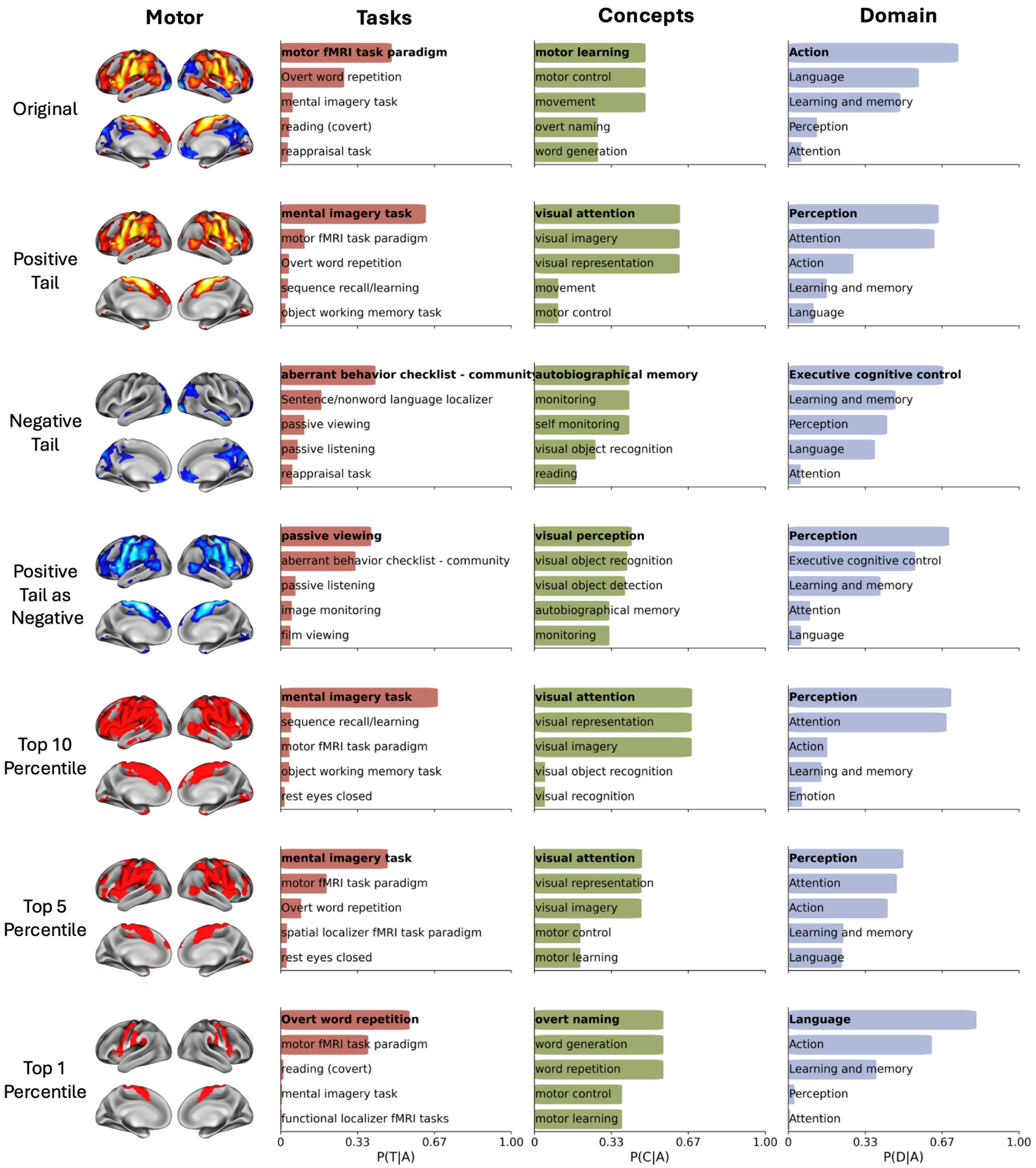


**Figure S9. NiCLIP for predicting tasks, concepts, and domains from different variations of the same motor group-level maps.**

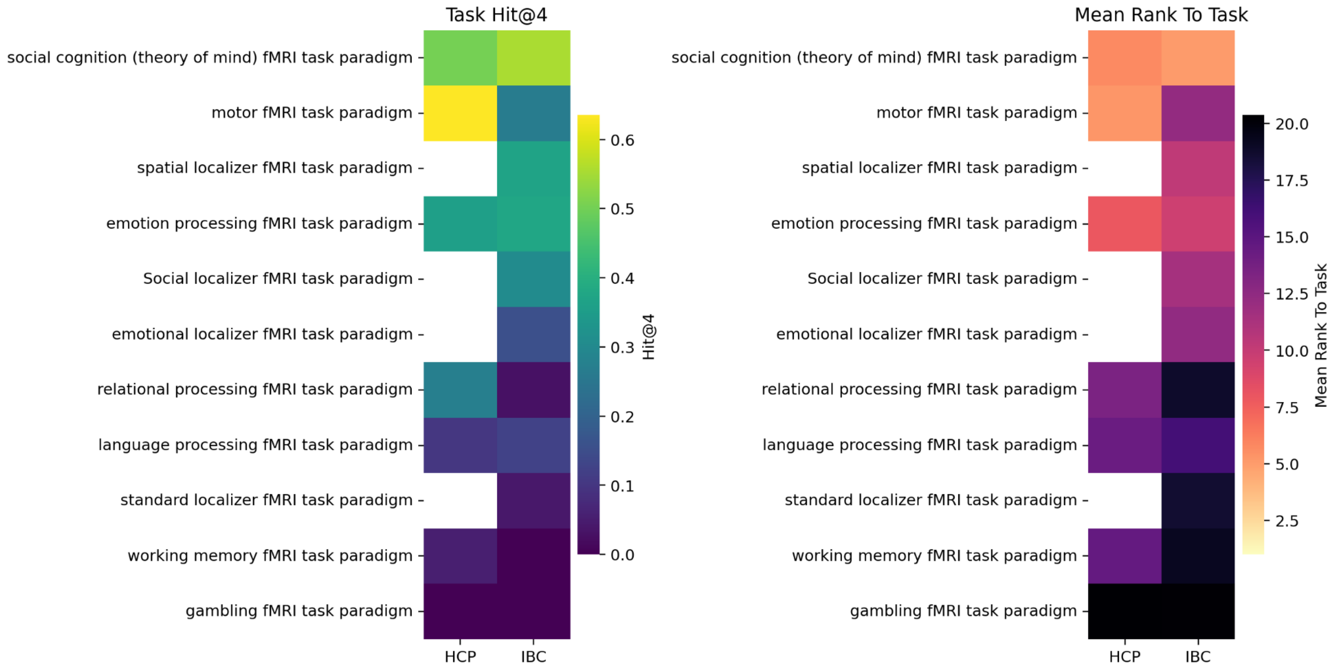


**Fig S10. Task-level decoding accuracy across HCP and IBC datasets.** Decoding was evaluated by retrieving the top-ranked terms from the full Cognitive Atlas vocabulary for each brain image. (Left) Hit@4: proportion of images for which the correct task term appeared in the top 4 retrieved terms. (Right) Mean rank of the correct task term across all images. Tasks are ordered by mean Hit@4 across datasets. Social cognition and motor tasks are decoded most reliably, while gambling shows the weakest performance, consistent across both datasets.

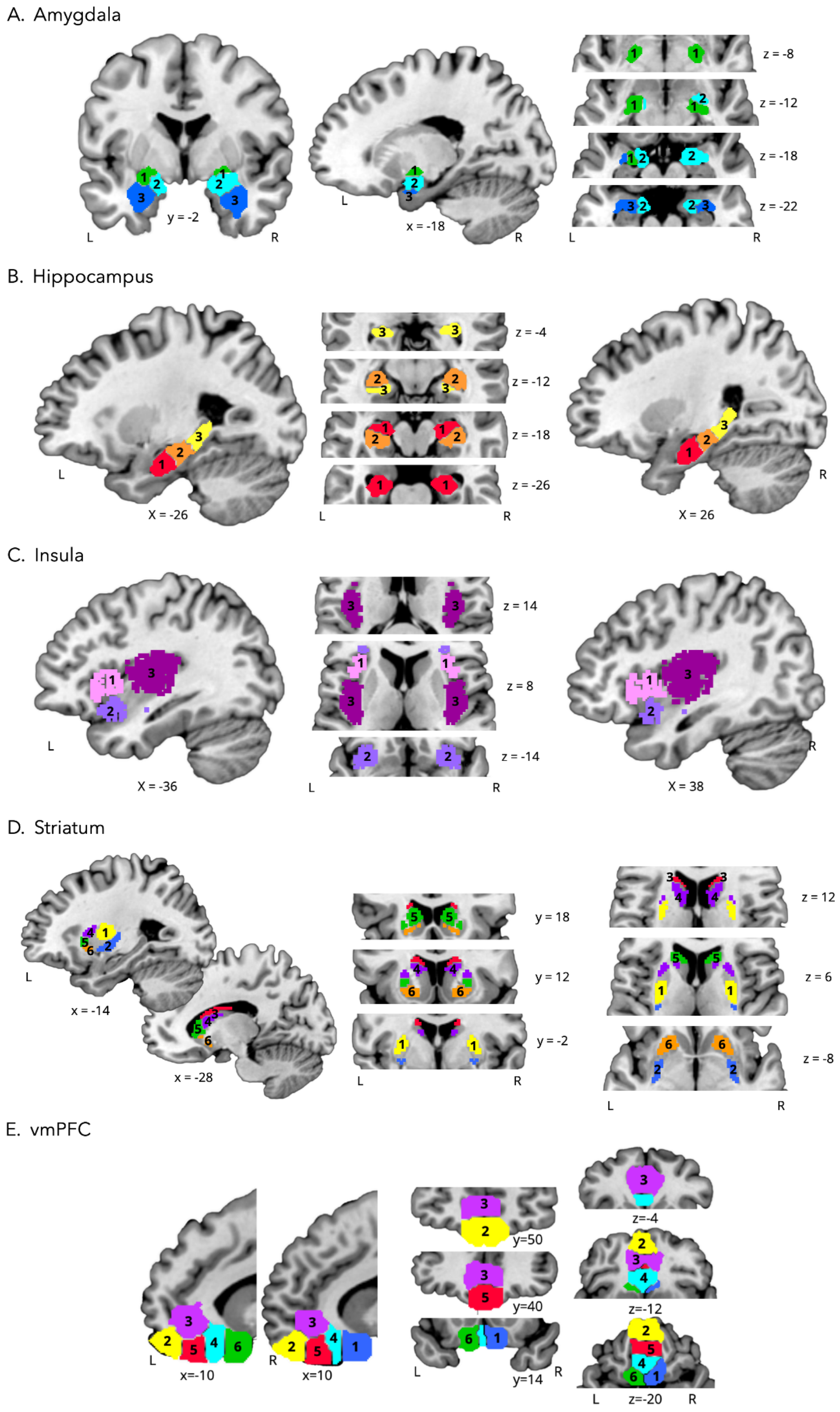


**Figure S11. Meta-analytic regions of interest.**
